# ATP-Binding Free Energy Simulations Reveal an Allosteric Link Between the Enzyme Active Site and Multiple Functional Protein-Protein Interaction Interfaces in Cyclin Dependent Kinase-1

**DOI:** 10.1101/2024.01.15.575662

**Authors:** Krishna Kant Vishwakarma, Ullas Seetharam Kolthur, Ravindra Venkatramani

## Abstract

The ATP dependent phosphorylation activity of Cyclin Dependent Kinase 1 (CDK1), an essential enzyme for cell cycle progression, is intrinsically dependent upon interactions with Cyclin-B, substrate, and Cks proteins. A recent joint experimental-computational study from our group showed intriguingly that acetylation at the active site abrogated the binding of CDK with Cyclin-B. These results posit the possibility of a bi-directional communication between the catalytic site and the protein-protein interface(s). Now, we present evidence for a general allosteric link between the CDK1 active site and all three of its protein-protein interaction (PPI) interfaces through atomistic molecular dynamics simulations (MD). Specifically, we examined ATP binding free energies to CDK1 in native non-acetylated (K33wt) and acetylated (K33Ac) forms as well as in two mutant forms of CDK1, the acetyl-mimic K33Q and the acetyl-null K33R which are accessible in-vitro. We find, in agreement with experiments, that ATP binding is energetically more favorable in K33wt relative to all other states wherein the active site lysine is perturbed (K33Ac, K33Q, and K33R). We also develop an entropy decomposition scheme which reveals, in addition to expected local changes in entropy, significant non-local entropy responses to ATP binding/perturbation of K33 from the 𝛼𝐶-helix, activation loop (A-loop), and the 𝛼𝐺-𝛼H loop segments in CDK1 which interface with Cyclin-B, substrate, and Cks proteins. Statistical analyses over a large set of MD trajectories reveal that while the local and non-local entropic responses to active site perturbations are on average correlated with dynamical changes in the associated protein segments, such correlations may be lost in about 9-48 % of the dataset depending on the segment. Besides, proving the bi-directional communication between the active site and CDK1:Cyclin-B interface, our study provides new insights into the regulation of ATP binding by multiple PPI interfaces in CDK1.

## 1. Introduction

Cyclin-dependent kinases (CDKs) are a family of proteins which serve as molecular regulators of the cell cycle in eukaryotes^1–3^. Dysregulation of CDKs results in abnormal proliferation of cells and diseases such as cancer.^4–8^ At present the prevalent view of activation of specific CDKs during various phases of the cell cycle is based on the temporally controlled expression of their cognate partner Cyclin proteins. The CDK enzymes are consistently expressed throughout the cell cycle and exist in an inactive state by default. The inactive CDK state is characterized by the presence of inhibitory post-translational modifications (PTMs) and an open active site. The activation of CDK involves the removal of the inhibitory PTMs, which include two phosphorylations at conserved threonine and tyrosine residues and an acetylation of an active site lysine. Subsequently, the binding of cognate Cyclin partners to the CDK N-lobe leads to a conformational change which closes the CDK active site and promotes phosphorylation of a conserved threonine in the activation segment, the hallmarks of a catalytically competent CDK:Cyclin complex^9–12^. Thus, the tight temporally controlled synthesis and degradation of cyclin proteins regulates CDK activity to drive the cell cycle. Additionally, the activity of the CDK:Cyclin complex is further tuned temporally by CDK inhibitors (CKIs) and CDK subunit (Cks) proteins. For instance, CKIs inhibit CDK activity by binding to either free CDK or to CDK:Cyclin complexes to inhibit either Cyclin or ATP binding respectively ^13–16^. In contrast, the Cks proteins bind to the C-lobe of CDK, distinct from the Cyclin binding site (**Fig. 1a**) and interact with substrate proteins to increase their specificity to CDK ^12,17^ In the case of the two inhibitory PTMs, the phosphorylation of the G-loop in CDKs is known to hinder substrate binding and to create an unproductive ATP conformation ^18–20^. However, the molecular mechanisms by which acetylation at the highly conserved active site lysine^21^ regulates CDK function are not completely understood. Acetylation of the catalytic lysine in CDK2, CDK5, and CDK9 has been shown to lead to a loss of enzyme function ^22–24^. Acetyl-mimic mutations changing the catalytic lysine to glutamine also result in a complete loss of enzyme function ^21^. Note that while the lysine to glutamine mutation mimics the natural PTM (acetylation) in terms of eliminating the charge, it does not preserve residue shape complementarity. Interestingly, the acetyl-null lysine to arginine mutation which preserves the charge of the residue, but adds a bulkier side-chain also leads to a loss of enzyme function ^21^. In CDK5 and CDK9, the loss of enzymatic activity upon acetylation was attributed to impaired ATP binding ^22,24^. These results suggest that the effects of acetylation are local and restricted to the active site of the CDK proteins.

**Figure 1:**
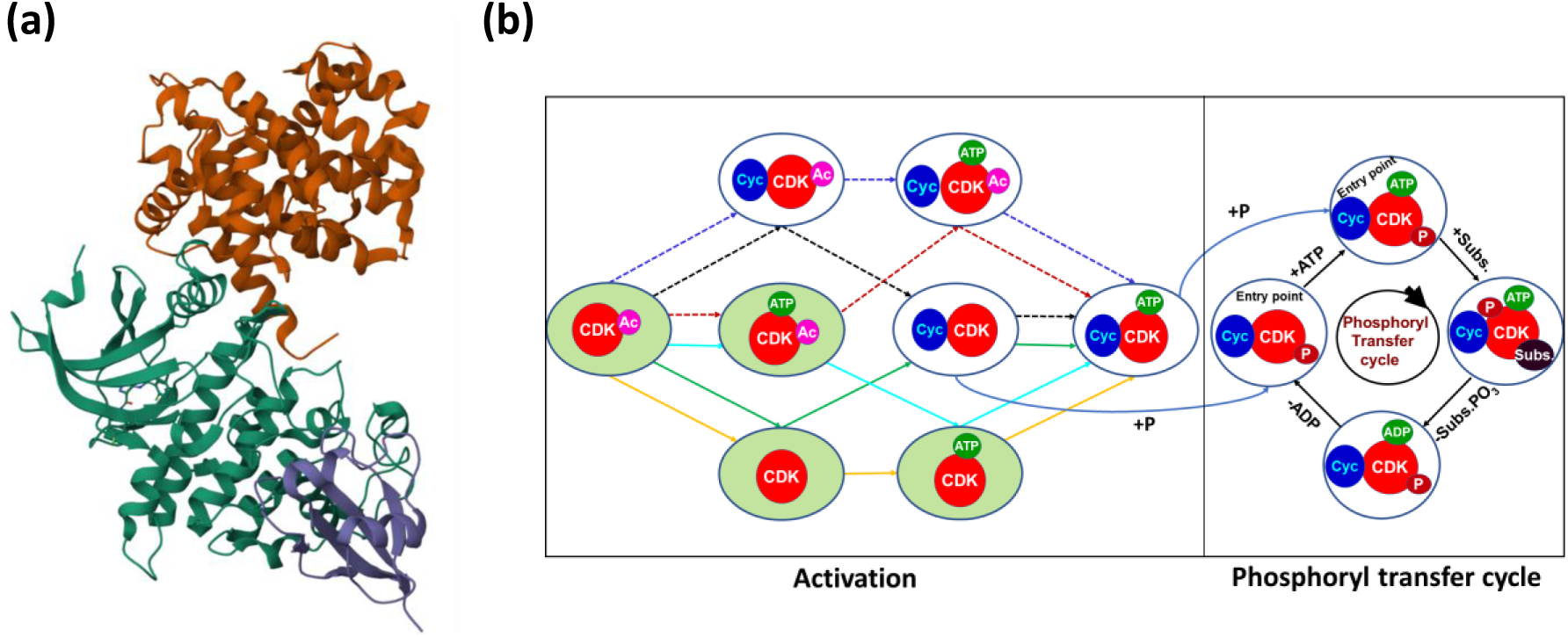
**(a)** Structure (PDBid: 4Y72) of CDK1 (green) bound to Cyclin-B (orange) and Cks2 (blue), rendered using Mol* 3D viewer from rscb.org **(b)** Simplified schematic representation of possible activation pathways for acetylated CDK1 leading up to the phosphoryl transfer cycle of an activated CDK1:Cylin-B complex. The simplified scheme does not include the steps involving the regulatory CKIs and Cks proteins and inhibitory phosphorylation steps. Favourable and unfavourable pathways, as determined by ref 25 are represented by solid and dotted arrows respectively. The four states of Cyclin free CDK1 lying in the green shaded circles are studied here.

Recently, we investigated the role of the catalytic lysine (K33) and its acetylation on CDK1 function^25^. Among the four CDKs which orchestrate cell cycle progression (CDK1, CDK2, CDK4, and CDK6), CDK1 is the only essential kinase driving the G2-M transition and mitosis. Using both biochemical assays and computational molecular dynamics (MD) simulations, we extracted molecular level insights on the loss of function in the CDK1:Cyclin-B complex upon perturbing K33; either by acetylation (K33Ac) or mutations (K33Q and K33R). We found that while the charged state of K33 is not essential for ATP binding, it nevertheless contributes to binding energetics and influences ATP conformations at the active site. More surprisingly, our studies revealed a non-local impact of K33 modifications/mutations on the Cyclin-B interface resulting in a significant reduction in Cyclin-B binding for the K33Q mutant relative to non-acetylated (K33wt) enzyme. Our study uncovered a previously unknown role for acetylation in regulating CDK1:Cyclin-B interactions. As shown in **Fig. 1b**, the acetylated CDK1 undergoes multiple activation steps including deacetylation at K33, ATP binding, Cyclin binding, and activatory phosphorylation before reaching a catalysis competent state. Thus, our previous study raises interesting mechanistic questions of the states of CDK1 where the loss of enzyme function observed due to K33 acetylations or mutations originates. Furthermore, it is not clear whether and to what extent uncomplexed (Cyclin-free) CDK1 contributes to the differences in ATP binding affinities for the K33Q/K33R mutants relative to K33wt observed in our biochemical assays^25^. Here, we address both of these issues by investigating free energies changes associated with ATP binding and K33 perturbations (via acetylation and K33Q/K33R mutations) on free CDK1 states along favourable enzyme activation pathways (green shaded circles of **Fig. 1b**). A detailed analysis uncovers a surprising non-local communication between the active site and segments of CDK1 which interface with Cyclin-B, substrate and Cks proteins. The results present molecular mechanisms by which CDK1 might regulate the binding of Cyclin-B and also sheds light on how the loss of enzyme function might arise through related but distinct perturbations of the catalytic lysine site.

## 2. Methods

### 2.1. Modelling of solvated CDK1 systems

Two sets of free CDK1 model systems (minus Cyclin-B and inhibitory phosphorylations) were prepared with and without ATP. All systems were derived from a crystal structure for CDK1 bound to Cks protein reported by Endicott and co-workers (PDB ID: 4YC6)^12^. The reported CDK1 structure contained missing residues within the protein chain (residue ID: 155-158) and at the C-terminus (residue ID: 290-297). We removed the Cks protein and added non-terminal missing residues in CDK1 using Modeller (version 9.19) ^26^ and hydrogen atoms using VMD^27^. The structure of the CDK1:Cyclin-B complex (PDB ID: 4Y72) also reported by Endicott and co-workers ^12^ was used as a template to model the missing residues (155-158) within the protein chain. The resulting structure was then solvated in a cubic water box (TIP3P water model) of size ∼93×93×93 Å^3^ with a minimum protein to box edge separation of 10 Å. The protein sequence has a net charge of +1 at pH 7 and a single Cl^-^ ion was added to arrive at a solvated and neutral ATP free non-acetylated native system (K33wt). Three perturbed solvated models without ATP were further derived from K33wt where the catalytic lysine was changed to glutamine (K33Q), acetyl-lysine (K33Ac), and arginine (K33R). For K33Q and K33Ac no Cl^-^ counterions were required as the system is neutral at pH 7. Subsequently, for each of these structures, we docked ATP at the active site of the enzyme to create four more systems: K33wt:ATP, K33Ac:ATP, K33Q:ATP, and K33R:ATP. Here, to dock ATP in the active site pocket of each CDK1 structure, we used a CDK2 crystal structure bound to ATP (PDB ID: 1B38) as a reference^28^. The reference and target CDK1 structures were aligned using VMD MultiSeq^29^and the resulting coordinates of ATP from CDK2 were assigned to the CDK1 active site. The conformation of the ATP in the active site pocket was refined during molecular dynamics equilibrations (*vide infra*). The complete list of systems thus generated is provided in **Table S1** of ESI.

### 2.2. Molecular Dynamics Simulations

We employed GROMACS (version 2020.1)^30^ to carry out atomistic MD simulations of each solvated CDK1 model using the CHARMM36^31^ force field and TIP3P^32^ water model. We used periodic boundary conditions along with the Particle Mesh Ewald (PME) method to compute full electrostatics. Further non-bonded interactions were truncated at 12 Å using Verlet^33^ cut-off scheme and a switching function (force-switch) for van der Waals interactions at 10 Å. Each model system was subjected to an equilibration protocol comprising multiple stages of minimization, heating, and pressure equilibration as follows. In the first stage of minimization and heating equilibration, the protein heavy atoms (and ATP and Mg^2+^ ions if present) were restrained with a force constant of 1000 kJmol^-1^nm^-2^ while the protein hydrogen atoms, water, and solvated ions (Na^+^/Cl^-^) were unconstrained. The systems were minimized using the steepest descent algorithm till the maximum forces on the system became less than 1000 kJmol^-1^nm^-1^. Then the systems were heated from 0 to 303 K in single step and equilibrated at 303 K for 125 ps using Nosé-Hoover thermostat^34^. In the second stage of minimization and heating equilibration, the steps from the first stage were repeated after removing the constraints on modelled/mutated residues. Additionally, there were no constraints on protein hydrogen atoms, water, and solvated ions. The third stage of minimization and heating equilibration was identical to the second stage except for the value of force constant (reduced to 100 kJmol^-1^nm^-2^). After these steps, systems were subjected to four rounds of NPT equilibrations at a temperature and pressure of 303 K and 1 bar respectively with successively lower constraints on protein heavy atoms (force constants of 100, 10, 1, and 0.1 kJ mol^-1^nm^-2^) using the Nose-Hoover thermostat and the Berendsen barostat^35^. Here, all atoms from mutated/modelled residues remained unconstrained. Finally, each model system was subjected to 60 independent unconstrained NPT simulation runs of 1 ns under the same condition as the previous NPT steps. Following equilibration, in the production phase, 60 independent 50 ns long MD trajectories for each system were generated under NPT conditions from the last set of coordinates from the previous step using the Parinello-Rahman^36^ barostat and the Nosé-Hoover thermostat. The total number of production simulations and their parameters are summarized in Table S1 of ESI. Structures for the eight CDK1 systems generated after NPT equilibration (before the unconstrained 1 ns NPT runs) are shown in **Fig. S1** of ESI.

### 2.3. Free energy calculations

We estimated the ATP binding free energies in all four CDK1 systems as

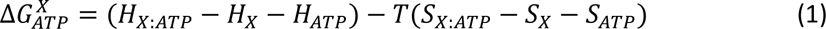

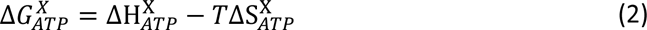

where X is K33wt/Ac/Q/R. Further, the relative free energy difference between K33wt and the corresponding unperturbed systems can also be estimated as

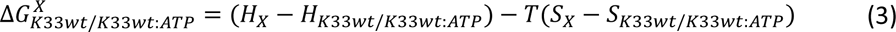

Or

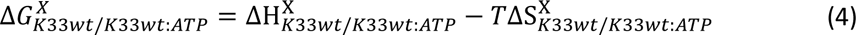

where *X* is K33Ac/Q/R or K33Ac/Q/R:ATP for the ATP free or ATP bound case. Here, the enthalpic part of free energy (𝐻) was estimated by Molecular Mechanics Generalized Born Surface Area (MM-GBSA) method in NAMD^33,37^. In the MM-GBSA method, the enthalpy 𝐻 is written as

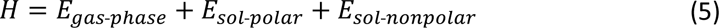

Here the 𝐸_𝑔𝑎𝑠-𝑝ℎ𝑎𝑠𝑒_ is the gas-phase energy of the molecule which is approximated using the molecular mechanics force-field^37^. 𝐸_𝑠𝑜𝑙-𝑝𝑜𝑙𝑎𝑟_ and 𝐸_𝑠𝑜𝑙-𝑛𝑜𝑛𝑝𝑜𝑙𝑎𝑟_ are the polar and non-polar parts of the solvation energy respectively. Here the polar part is calculated using the generalized Born method^38^, whereas the non-polar part is estimated from the solvent-accessible surface area (SASA) assuming a surface tension (proportionality constant) value of .005 kcal/mol/Å^2^. For each 50 ns trajectory, a total of 250 frames (a sampling rate of 200 ps) were used to calculate the enthalpy of the CDK1 system. For enthalpy calculations, we considered the protein, ATP and Mg^2+^ ions with active site waters. The latter were defined for every snapshot in the trajectory as the 50 most proximal water molecules as defined by the distance between their oxygen atoms and the 𝐶_𝛼_ atom of residue D146. The free energy change of ATP binding as a function of number of water molecules around D146 converges with 50 waters (**Fig. S2**). For the entropic contribution to free energy, we considered the vibrational, rotational and translations contributions. The vibrational contributions were estimated using the quasi-harmonic prescription provided by Andricioaei and Karplus ^39^ as

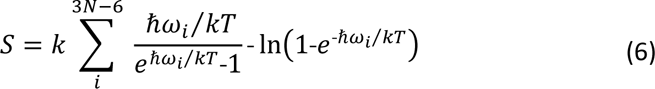

Here 𝑘 is the Boltzmann constant, 𝑁 is the total number of atoms, 𝑇 is the temperature, and 𝜔_𝑖_is the mode frequency calculated from a principal component analysis (PCA) of the atomic fluctuations from MD trajectories. PCA involves the diagonalization of a mass-weighted covariance matrix 𝐶 (elements 𝐶_𝑖𝑗_ = 〈𝑥_𝑖_ − 〈𝑥_𝑖_〉〉 〈𝑥_𝑗_ − 〈𝑥_𝑗_〉〉) of atomic coordinates 𝑥_𝑖/𝑗_(*i,j*=*1…N*) constructed from an *N*-atom MD trajectory, free of rigid body rotations and translations, to yield 3*N*-6 uncorrelated modes (eigenvectors 𝜉_𝑚_) and their corresponding frequencies 𝜔_𝑚_(eigenvalues 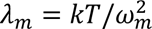). The covariance matrix was generated using MD coordinate frames at a sampling rate of 2ps. The sampling rate for entropy calculations is much higher than that for enthalpy (200 ps) as the former was found to be strongly dependent on sampling and produces small error of mean at high sampling rates (**Table S2**). On the other hand, the average enthalpy does not change significantly with sampling rate (**Table S2**).

For our analysis, we extracted the variance contributions from specific subsets (protein segments) by tracing over the variance contributions 𝐶_𝑖𝑖_ from atomic coordinates 𝑥_𝑖_ that belong to that subset. Changes in variance for a specific protein segment (*p*) with ATP binding or upon perturbing the active site lysine are then estimated as:

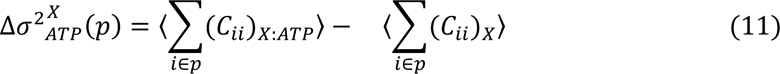

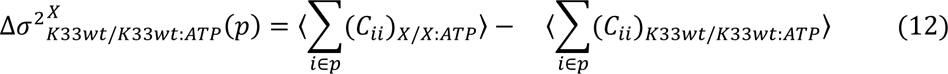

Where the averages are over the 60 sets of 50 ns trajectories. The transformation of the entropy 𝑆 matrix elements in the mode basis (𝑆_𝑚𝑛_) to the atomic basis (𝑆_𝑖𝑗_) is obtained from the eigenvectors 𝜉_𝑚_of 𝐶 as:

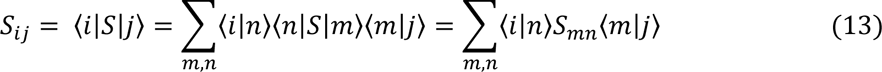

Since 𝑆_𝑚𝑛_ = 0 for 𝑚 ≠ 𝑛

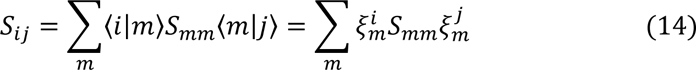

where 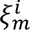 is the expansion co-efficient of the 𝑖^𝑡ℎ^ atomic coordinate for the *m*^th^ eigenvector 𝜉_𝑚_. Therefore, the contribution of the *i*^th^ atomic coordinate to the entropy is:

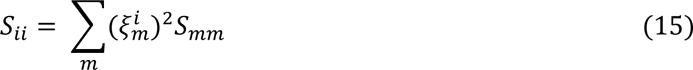

The entropy contributions from specific subsets (atoms, residues, or segments) is then simply given by tracing over the entropy contributions from coordinates that belong to that subset. Changes in entropy contributions for a specific protein segment (*p*) with ATP binding or upon perturbing the active site lysine are then estimated as:

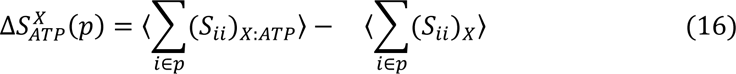

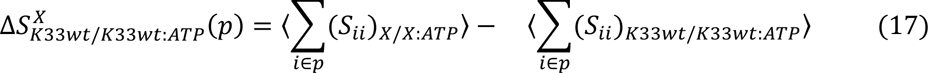

where the averages are over the 60 sets of 50ns trajectories.

For translational and rotational entropy, we have used the approach of ideal gas entropy terms at 1 M concentration as suggested in reference ^40^. The expression for translation and rotation entropy of molecule at concentration 𝐶^0^= 1M is given by

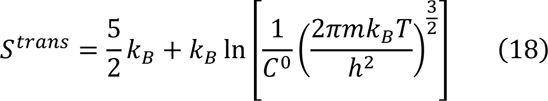

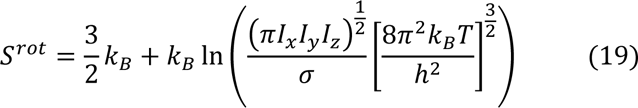

Here ℎ is the Planck’s constant, m is the mass of the molecules, 𝐼_𝑥_, 𝐼_𝑦_, 𝐼_𝑧_ are the moment of inertia of the molecule, and 𝜎 is the symmetry factor whose value is taken 1 for this work.

## 3. Results

### 3.1. Perturbation of the active site lysine in free CDK1 reduces its ATP binding affinity

Several studies, including our previous investigation of CDK1 have showed that the positive charge of K33 is dispensable for ATP binding in kinases ^25,41,42^. In our studies on CDK1, ATP competition assays and binding to ATP analogs revealed ATP binding to be slightly reduced in K33Q and K33R mutants relative to the unperturbed system^25^. However, it remained unclear to what extent free CDK1 accounts for the reduced ATP binding. Here, we find, in agreement with experimental observations, that the free energy of ATP binding in CDK1 without Cyclin-B indeed reduces by 36, 40 and 12 kcal/mol, respectively when acetylating K33 or mutating it to either glutamine or arginine, respectively, although binding still remains favourable in these perturbed systems (**Fig. 2a**). A decomposition of free energy changes into enthalpy (*ΔH*) and entropy (-*TΔS*) contributions reveals that ATP binding induces favourable enthalpy changes and unfavourable entropy changes in all systems which counterbalance to produce the observed trends. Abrogation of the positive charge on K33 significantly reduces the quasiharmonic entropy contributions associated with conformational changes. We find the largest energetic changes (both enthalpy and entropy) for the charged systems (K33wt and K33R). These systems show a more favourable enthalpy and unfavourable entropy change upon ATP binding than the neutral K33Q/K33Ac mutants. This result is along expected lines as the active site should respond more strongly to the ATP when the K33 carries a charge. Between K33wt and K33R, the former shows a larger favourable enthalpic change and the latter shows a more unfavourable entropy change (**Fig. 2b**). A decomposition of enthalpy changes into bonded/non-bonded interactions and solvation energy reveals large opposing nonbonded and solvation energy contributions (**Table S3**). In all cases, modifications of the active site K33 leads to unfavourable electrostatic and vdW energy changes (∼100-165 kcal/mol) which are countered by smaller but favourable solvation energy contributions (∼60-120 Kcal/mol) relative to K33wt. Interestingly, the perturbations which abrogate the charge on K33 produce a larger unfavourable vdW change in the *Δ**H*** for ATP binding relative to K33R which preserves the charge. However, the trends for the change in electrostatics upon perturbing K33 is not so consistent. The K33Q mutation, which replaces the ATP-tail coordinating lysine with a smaller uncharged residue does induce the largest unfavourable change in electrostatics, almost 40 kcal/mol higher than that for K33R, as intuitively expected. However, surprisingly, the similarly positive to neutral K33Ac change induces a much lower (by 20 kcal/mol) unfavourable electrostatics than the K33R mutation which preserves the charge. In order to get further insights into active site residue specific contributions to the non-bonded enthalpy change with ATP binding, we examined changes in ATP (carrying Mg^2+^) electrostatic and vdW interaction energies with active site residues (**Fig S3**). Significant non-bonded interactions (dominated by electrostatics) of the ATP in K33wt in descending order of strength is with K130, K33, D146, D86, N133, T14, G12, and G13. Here, only the interaction with D86 is repulsive which partially offsets the strongly attractive interaction with K130. The perturbations at K33 only impact the K33 site and the T14 (phosphorylation) site weakening their interactions with ATP. In the K33Q/Ac wherein the charge on K33 is eliminated the impact is the greatest. Solvation energy compensates in these systems to keep ATP binding favourable and competitive relative to that in K33R.

**Figure 2:**
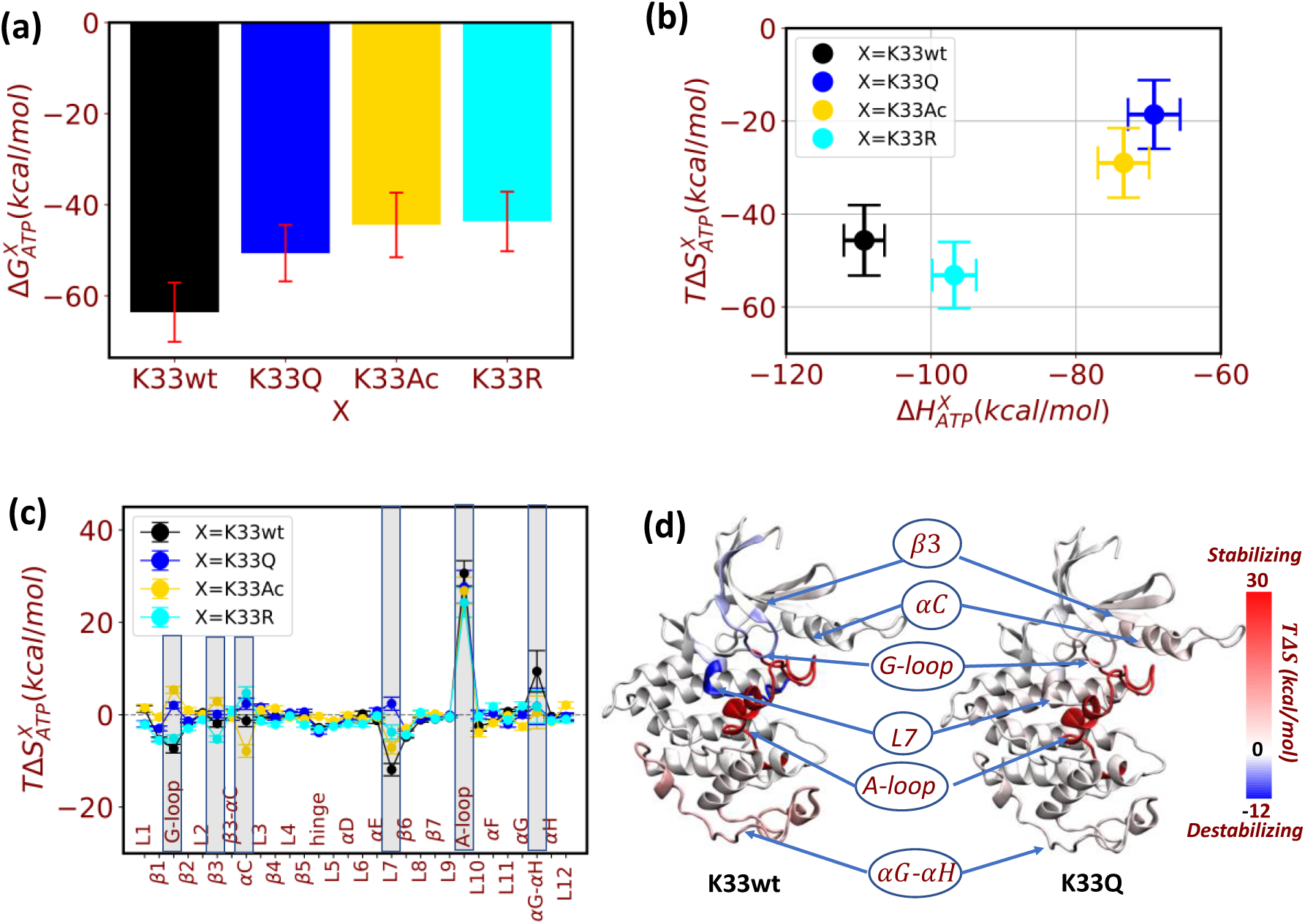
Effect of mutations/modifications at K33 on ATP binding in CDK1. **(a)** Changes in free energy 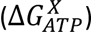, of ATP binding in CDK1 **(b)** Correlation of enthalpy 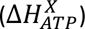 and entropy 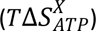 changes in CDK1 associated with ATP binding. Here, averages and standard errors of the mean were extracted from 60×50 ns trajectories for each system (X=K33wt, K33AC, K33Q and K33R). **(c)** Contributions to 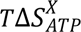 from distinct segments of CDK1. Prominent entropic changes in CDK1 with ATP binding are indicated by the shaded boxes. **(d)** Heat maps showing the entropy changes upon ATP binding in the protein structure for K33wt and K33Q. Heat maps for other systems are provided in the ESI Fig. S7.

We find that the inclusion of active site water molecules is essential to produce the correct trends in ATP binding free energies (compare **Fig 2a**. and **Fig S4a)**. When active site waters are excluded, not only is ATP binding unfavourable for all systems (*Δ**G** > 0*), but the K33wt system shows comparable or slightly worse affinity for ATP than the neutral systems. This trend is both non-intuitive and in disagreement with experiments ^25^. While the enthalpy change with ATP binding is still favourable in K33wt, the value of *Δ**H*** is significantly reduced so that the free energy trends are now dominated by the entropy change. When explicit active site waters are included, *Δ**H*** for ATP binding in K33wt also become more favourable relative to K33Q/Ac (**Fig S4b** and **2b**). We find that the inclusion of active site waters predominantly destabilizes the electrostatic interactions and stabilizes the solvation energy contributions selectively in the ATP-bound states of the K33Q/Ac systems to accentuate the *Δ**H*** differences of these systems relative to K33wt (**Tables S3-S5**). In contrast, presence of active site water molecules only slightly impacts the relative difference in *Δ**H*** between K33wt and K33R (**Fig S4b** and **2b**). From these observations we can conclude that nullifying the charge on K33 significantly changes the interactions (unfavourably) between ATP and waters at the active site of the enzyme.

### 3.2. Local and non-local entropic contributions to ATP binding free energies in native and mutant CDK1

The data in **Fig. 2b** indicates that binding of ATP in K33wt leads to significant unfavourable entropy changes which are much larger than that seen in neutral mutant systems (K33Q/Ac) but comparable to that in the charge preserving K33R. In our calculations we account for vibrational, translation and rotational entropies. However, the four CDK1 systems differ only in terms of their vibrational entropy as the translational and rotational entropies provide identical contributions to ATP binding free energies in our model (**Table S6**). The binding of a ligand to an enzyme active site is expected to locally decrease the flexibility, leading to an unfavourable entropy change. The correlation between flexibility and entropy will be explored in more detail later in **subsection 3.4**. ATP binding in K3wt/R decreases its vibrational entropy as expected. However, ATP binding to K33Q/Ac does not follow expectation and increases its the vibrational entropy (**Fig S5a**). To investigate these trends, we decomposed the entropy changes upon ATP binding (Δ𝑆_ATP_^𝑋^) in all four CDK1 systems into per residue contributions (**Eqn.16**). In **Fig. S5b**, we plot the total entropy change arising from residues as a function of their distance from the D146-𝐶_𝛼_atom at the active site. The charged systems K33wt/K33R show unfavourable entropy changes with ATP binding not only near the active site as expected but surprisingly also from regions far away from the active site (> 10 Å away from D146-𝐶_𝛼_). For the charge neutralizing mutants K33Q/K33Ac also large non-local favourable contributions dominate the entropy changes with ATP binding. Next, we decomposed the CDK1 sequence into 30 non-overlapping segments (**Fig. S6 and Table S7**) comprising of helices, strands and loops and computed their contributions to the overall entropy change for each system (**Fig. 2c**). These calculations reveal that the largest entropic changes created by ATP binding for the four CDK1 systems arise from local (interacting with ATP) contributions from the L7, 3, and G-loop segments and non-local contributions from the activation loop (A-loop), 𝛼𝐶-helix and the 𝛼𝐺-𝛼H-loop (**Fig. 2c-d and S7**). As can be seen from **Table S7**, the residues G13 and G14 belong to G-loop, K33 is placed within 3, and K130 is associated with L7. These residues form favourable non-bonded interactions with ATP (**Fig S3**) suggesting expected change in entropy from these segments upon ATP binding. ATP binding is found to consistently decrease the entropy contributions of the local regions (L7, 3, and G-loop) in K33wt/R the changes are inconsistent for the K33Q/K33Ac systems. While the G-loop entropy non-intuitively increases for both K33Q/Ac, entropy of the L7 segment increases for K33Q and decreases for K33Ac, and the 3 entropy shows a slight increase for K33Ac and no significant change for K33Q. Non-locally, upon ATP binding to K33wt, the entropies of the A-loop and the 𝛼𝐺-𝛼H-loop increase to favour the ATP bound state (**Fig. 2c-d**). In contrast, the favourable entropy changes from these segments are either reduced to varying extents (𝐴-loop) or no changes (the 𝛼𝐺-𝛼H-loop) upon perturbing the active site lysine (**Fig. 2c-d and S7**). Intriguingly, the 𝛼𝐺 − 𝛼H loop and the A-loop interface with Cks and substrate protein respectively. Moreover, ATP binding in K33Ac also leads to significant changes in 𝛼𝐶-helix which binds to Cyclin-B, which is line with our observations of acetylation in CDK1 abrogating this binding interaction ^25^. In Akt kinase, ATP binding prevents the dephosphorylation of T308 (equivalent to T161 in CDK1) by reducing its exposure to phosphatases^43^. Here, we also find that the ATP binding alters the solvent accessibility of T161 (**Fig. S8**), albeit exposing the residue. Apart from ATP binding, the presence of Cyclin also impacts the conformation of the A-loop and solvent accessibility of T161 promoting an activatory phosphorylation at this site. Nevertheless, our results here clearly indicate the allosteric entropic contributions to ATP binding from functionally relevant PPI loop segments in CDK1.

### 3.3. Impact of modifying K33 on the ATP-free and ATP bound states of CDK1

Next, we compare the effect of mutations/acetylation at K33 on the free energies of CDK1 in ATP-free versus ATP-bound forms. The mutations which eliminate the charge on the active site lysine (K33Q and K33Ac) destabilize the ATP-free CDK1 relative to the K33wt counterpart (**Fig. 3a and Fig. S9a**). In contrast, the lysine charge preserving K33R mutation stabilizes the ATP-free CDK1 relative to ATP-free K33wt (**Fig. 3a and Fig. S9a**). In the absence of ATP, the K33Q and K33Ac mutations create to negligible enthalpic changes, and, non-intuitively, to significant unfavourable entropy changes thereby leading to a higher free energy of the mutant/acetylated systems relative to K33wt (**Fig. 3a and Fig. S9b-c**). For ATP-free K33R, both enthalpy and entropy changes are significant and favourable, stabilizing the mutant state relative to K33wt (**Fig. 3a and Fig. S9b-c**). ATP binding does not alter the relative trends in energetics between K33wt and K33Q/Ac/R (**Fig. 3a and Fig. S9a**). However, in the presence of ATP, the glutamine mutation or acetylation at K33 leads to unfavourable enthalpic changes while the arginine mutation leads to favourable change as in the ATP-free case (**Fig. 3a and Fig. S9b**). The K33Q/Ac:ATP systems exhibit favourable/unfavourable entropic changes, respectively, while K33R:ATP does not show a significantly change in entropy relative to K33wt:ATP (**Fig. 3a and Fig. S9c**). Upon decomposing the entropy changes into contributions associated with different segments, we again find for both, the ATP-free and ATP-bound case. prominent contributions from the local (L7, 3, and G-loop) and non-local (𝛼𝐶-helix, A-loop, and 𝛼𝐺-𝛼H-loop) regions (**Fig. 3b, S9d-i**). The extent to which each of these segments is impacted by a mutation at the active site as well as the sign of their entropy change are both sensitive to the specific mutation and the presence/absence of ATP at the active site. For instance, in both ATP-free and ATP-bound CDK1, acetylation at K33 results in a decrease in the entropy of the A-loop, while glutamine/arginine mutations increase the entropy of this segment. Mutations/acetylation in K33wt:ATP reduce the entropy of the 𝛼𝐺-𝛼H-loop, but do not change segment entropy significantly in the ATP free state. Mutations/acetylation in both ATP-bound and free states result in an increase and decrease in the entropy of 3 and the 𝛼𝐶-helix, respectively. Here, interestingly, the glutamine/arginine mutations selectively impact the ATP-free state, whereas acetylation selectively impacts the ATP-bound state. Moving on the local segments, almost all perturbations to K33 in the absence of ATP decrease the entropy of L7 and the G-loop but increase segment entropy when ATP is bound. The exceptions to this trend are the acetylation (L7) and the charge preserving K33R (G-loop) which in the absence of ATP do not significantly change entropy. While all perturbations (acetylation/mutation) of K33 in ATP-free CDK1 leads to entropy changes in two PPIs (𝛼𝐶-helix, A-loop), in the ATP-bound state of the enzyme the modifications at K33 impacts all three PPIs (𝛼𝐶-helix, A-loop, and 𝛼𝐺-𝛼H-loop). Taken together with the results of the last subsection and considering both ATP binding and mutations/acetylation at K33 as active site perturbations, it is apparent that the active site is allosterically coupled to three distinct regions of CDK1 (𝛼C-helix, A-loop, and 𝛼𝐺-𝛼H-loop) each of which are involved in forming interfaces with distinct protein partners. Furthermore, ATP binding seems to enhance the coupling of the K33 site to the 𝛼𝐺-𝛼H-loop (**Fig. 3b** and **S9d-i**).

**Figure 3:**
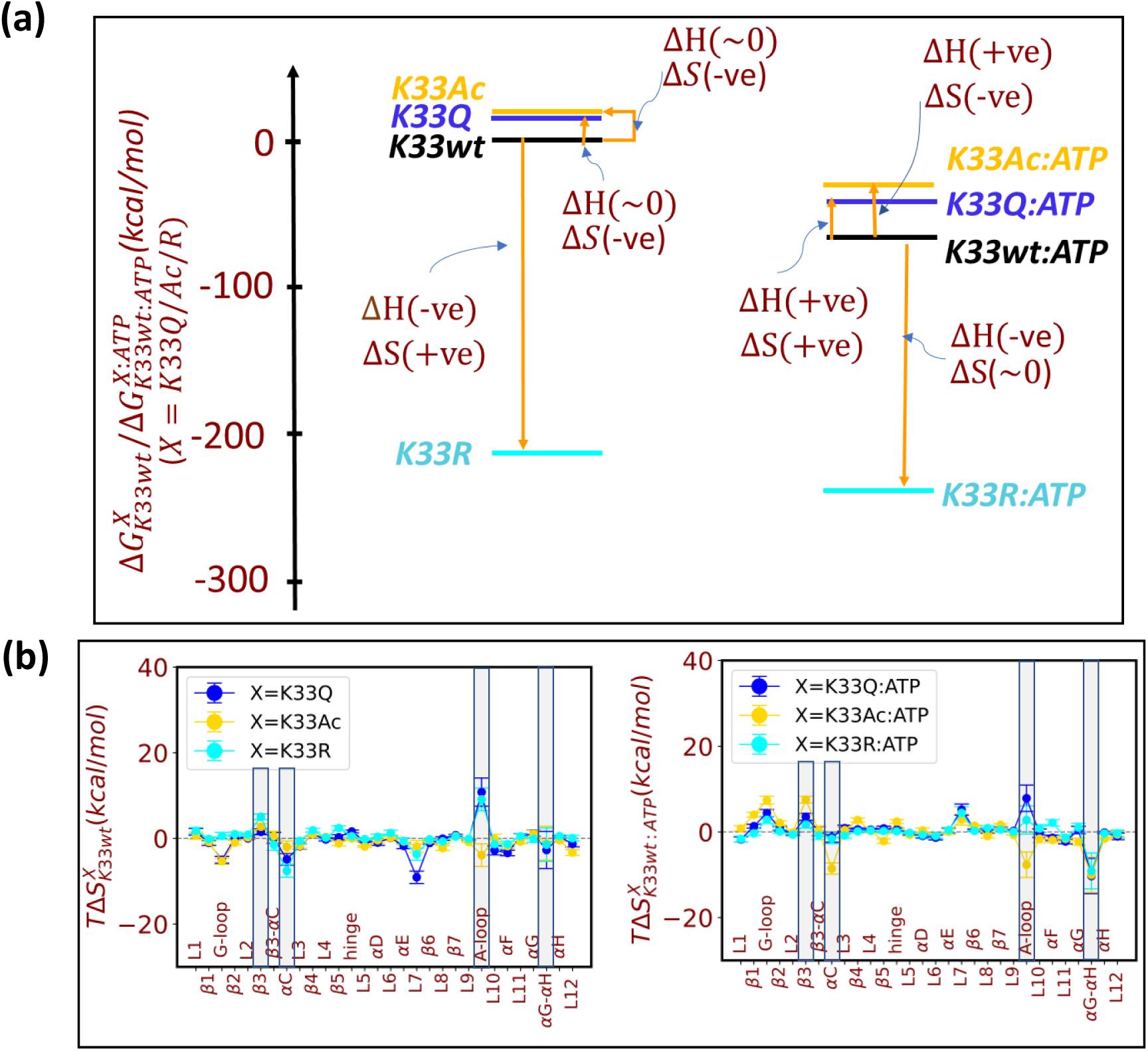
Effect of mutations/modifications at K33 on the free energies of ATP-free and ATP bound states of CDK1. **(a)** free energies of perturbed states of CDK1 in free form and when bound to ATP relative to K33wt (zero free energy). The relative free energy values reflect averages extracted from 60×50 ns trajectories for each system. We indicate the stabilization/destabilization of each perturbed ATP-free and ATP bound state in terms of enthalpy and entropy contributions relative to K33wt and K33wt:ATP respectively **(b)** Entropy change with mutation of K33 in 30 non-overlapping segments of CDK1 (see Fig. S6 and Table S7) for ATP-free (left subplot) and ATP-bound (right subplot) CDK1. The entropy changes for the A-loop, 𝛼𝐺-𝛼H-loop, and 𝛼𝐶-helix which interface with substrate, Cks, and Cyclin-B proteins are circled

### 3.4. Correlation between changes in entropy and CDK1 flexibility induced by active site perturbations

We estimated the changes in the variance (mean square fluctuations) of the 30 non-overlapping segments of CDK1 induced by ATP binding and mutations at K33 (**Eqns. 11** and **12**) in the 60 × 50 ns MD trajectories. We find that ATP binding at the active site in CDK1 (native/mutant) specifically triggers changes in the thermal fluctuations of the A-loop, 𝛼𝐺-𝛼H-loop, and 𝛼C-helix, PPIs which interface with substrate, Cks and Cyclin-B proteins respectively, and the G-loop and L7 segments, which locally interact with ATP (**Fig. 4a, S10a-c** (red data)). Conversely, mutations/acetylation at K33 in the ATP-free or ATP-bound CDK1 also trigger changes in the thermal fluctuations (red data in **Fig. S10d-i**) of the same segments depending on the nature of perturbation at K33 and presence and absence of ATP. These results reinforce the evidence for a coupling between the active site and the three key PPIs of CDK1 which was uncovered in **Subsection 3.2** by examining entropy changes (**Fig. 2c** and **3b**). Intuitively, increasing (decreasing) flexibility of a segment is expected to manifest in a higher favourable (unfavourable) entropy change. Cross-correlating the changes in segment entropies and thermal fluctuations upon ATP binding for our sets of 60 trajectories we find that these quantities are correlated as expected on average (**Fig4a, Fig 4b, and S10a-i**). However, two interesting observations can be made from the cross-correlation data. The first is that the relative extent of changes in thermal fluctuations between loop segments are not always commensurate with the corresponding entropy changes. For instance, upon ATP binding in K33wt, the entropy change of the A-loop is only 2.5 times more than that for L7 **(Fig 4a)** while the change in fluctuation for the former is ∼10 fold higher than the latter segment. Secondly, the correlation between entropy changes and changes in thermal fluctuations can break down for data from individual pairs of trajectories (e.g a specific pair of ATP-bound and ATP-free system trajectories). The existence of such cases is also apparent from the large amount of scatter seen in the cross-correlation plots for the individual loops (**Fig 4b**). For instance, in a specific K33wt and K33wt:ATP trajectory pair (**Fig 4b-c**), the 𝐴-loop exhibits an increase in flexible (positive change) which counterintuitively decreases its entropy (negative change) destabilizing the bound state. The 𝛼C-helix also exhibits a decrease in entropy with ATP binding which does not translate into flexibility change (**Fig 4b-c**). Furthermore while ATP binding significantly increases the flexibility of the 3-𝛼C loop this translate into very small change in entropy of 3-𝛼C loop (**Fig 4b-c**). Such inconsistent correlations between segment entropy and fluctuations are also seen in trajectory pairs of K33Q/Ac/R and K33Q/Ac/R:ATP, K33wt and K33Q/Ac/R, and K33wt:ATP and K33Q/Ac/R:ATP (**Fig. S11-12**). Our analysis indicates that across the various local and non-local segments here, 9-48 % of trajectory pairs show a lack of correlation between entropy changes and changes in thermal fluctuations (**Table S8**). These observations indicate that interpreting flexibility changes in terms of entropic contributions based on single (or a small number) trajectories can be misleading.

**Figure 4:**
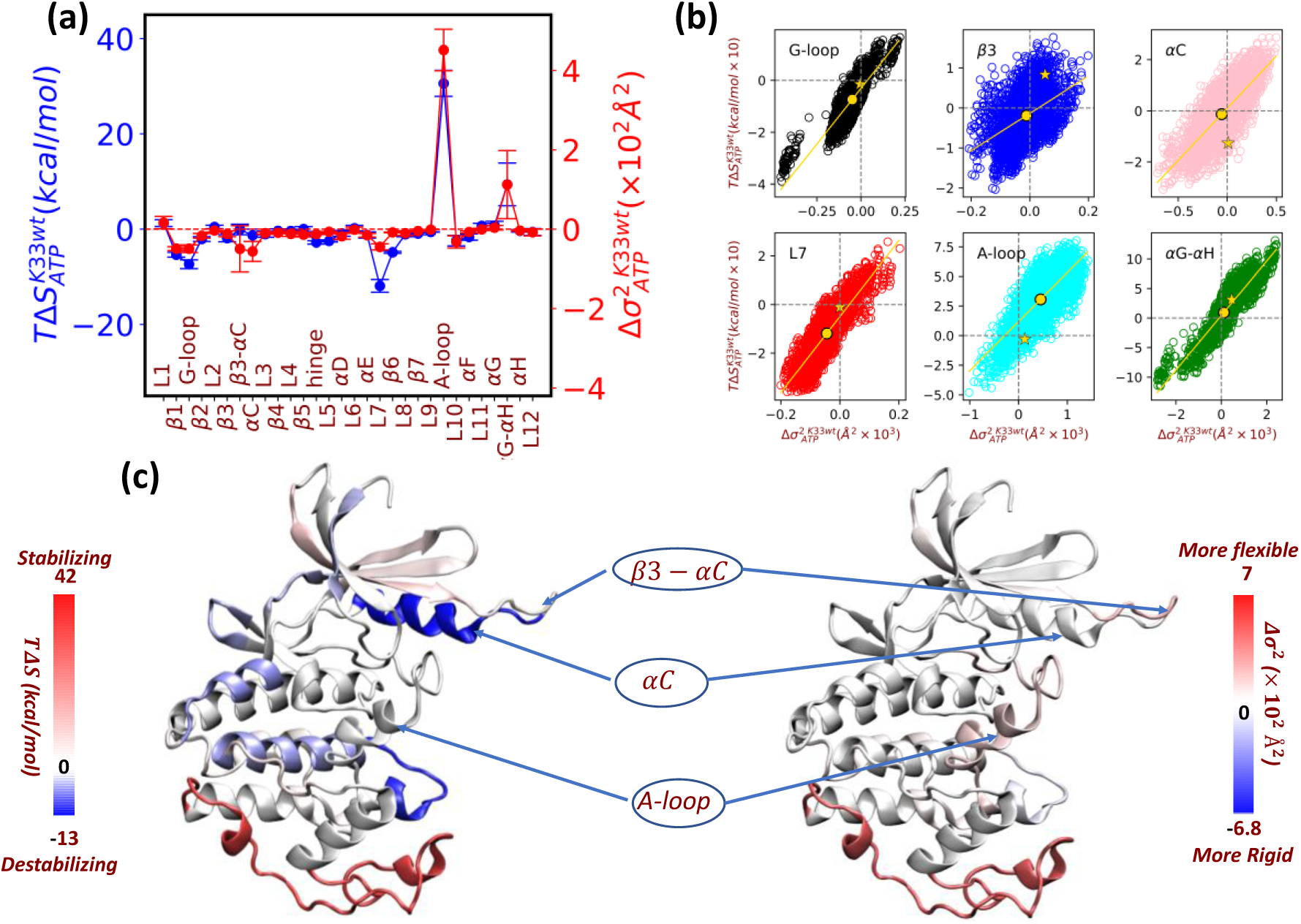
Correlation between the changes in the variance of atomic fluctuations and entropy with ATP binding to K33wt. **(a)** Entropy contributions to the free energies of ATP binding to K33wt (blue) from the 30 non-overlapping segments of CDK1 along with the corresponding changes in variance of atomic fluctuations (red). Data points reflect averages extracted from 60×50 ns trajectories for ATP-free and ATP-bound K33wt. Corresponding data for changes in the entropy and fluctuations of the segments with mutations at K33 are shown in ESI (Fig. S10a-c) **(b)** Scatter plot depicting the relationship between changes in the fluctuations (x-axis) and the entropic component of free energy (y-axis) of segments L7, β3, and G-loop which are local to the active site and non-local segments αC-helix, A-loop, and αG-αH-loop from 60 ATP free x 60 ATP-bound=3600 trajectory pairs for K33wT. Each unfilled circle on the plot represents a variance and entropy change from all possible pair of trajectories for these specific segments, each obtained from ATP-free and ATP-bound states. The filled circles indicate the averages for the segments which correspond to the data shown in (a). The filled star symbols indicate a representative trajectory pair for which the correlation between entropy change and changes in fluctuations are lost as indicated in the heat map plots shown in **(c).** The A-loop, β3-αC loop, and αC-helix are annotated here .

## 4. Discussion

In our previous study^25^, we showed for the first time that acetylation in CDK1 at the active site lysine (K33) regulates the binding between CDK1 and Cyclin-B. Our computational studies of a CDK1:Cyclin-B:ATP ternary complex revealed the mechanism of regulation to be an allosteric coupling between the active site of the kinase and the CDK1:Cyclin-B interface. Mutants/acetylation at the active-site lysine were found to slightly reduce the non-bonded interactions between the ATP and the enzyme albeit still keeping it favourable. However, a complete mechanistic understanding of the regulatory role of acetylation in CDK1 function requires an interrogation of ATP binding and mutations/acetylation at K33 in free CDK1 in the absence of Cyclin-B. Here, we have addressed this lacuna using free energy calculations and statistical analysis. Our studies reveal allosteric couplings between the active site and three PPIs, including with Cyclin-B, contained within the kinase architecture.

Our binding free energy calculations show that acetylation, acetyl-mimic, and acetyl-null mutations at K33 lower the ATP binding affinity, albeit keeping it favourable, in free CDK1. This is in accordance with experiments which show a reduced binding of FSBA to K33Q and K33R mutants^25^. Upon decomposing the free energy into enthalpy and entropy contributions, we find that ATP binding to K33wt predictably leads to favourable enthalpic changes localized at the active site. Intriguingly, however, there are also unexpected favourable entropic changes in the protein with large non-local contributions arising from two PPIs, the A-loop and the 𝛼𝐺-𝛼H-loop, which interface with substrate and Cks proteins. Further, in K33Ac states, ATP binding also leads to favourable entropic changes in the A-loop and a unfavourable entropic change at the 𝛼C-helix, which interfaces Cyclin-B protein. This surprising result indicates that the active site is coupled to multiple PPIs which can allosterically regulate the binding of ATP in free CDK1. We further find that mutations/acetylations at K33 in both ATP-free and ATP-bound CDK1 also produce significant entropy changes in three different PPI segments. The allosteric influence of the three distinct PPIs on the active site perturbations, including ATP binding, acetylation, and mutations at K33, is summarized in **Fig. 5**. Here, we examine the entropy change of the three PPIs for the 7 perturbed states (K33wt:ATP, X, and X:ATP where X= K33Ac, K33R, K33Q) relative to the unperturbed K33wt state of the enzyme. **Fig. 5** shows that the entropy of the Cyclin-B binding 𝛼C-helix in ATP-bound CDK1 is lowered to selectively disfavour acetylation. The 𝛼C-helix entropically also disfavours the glutamine and arginine mutations in ATP-free CDK1. In contrast, the substrate A-loop is influenced by almost all alterations to the K33wt state and shows significant increases in entropy to favour the perturbed states. The single exception is the acetylation of K33 in the absence of ATP which produces a small negative entropy change. Finally, the entropy of the Cks binding 𝛼𝐺-𝛼H segment increases to favour ATP binding to K33wt but is insensitive to all other perturbations. Taken together, all three PPIs entropically influence ATP binding to CDK1.

**Figure 5:**
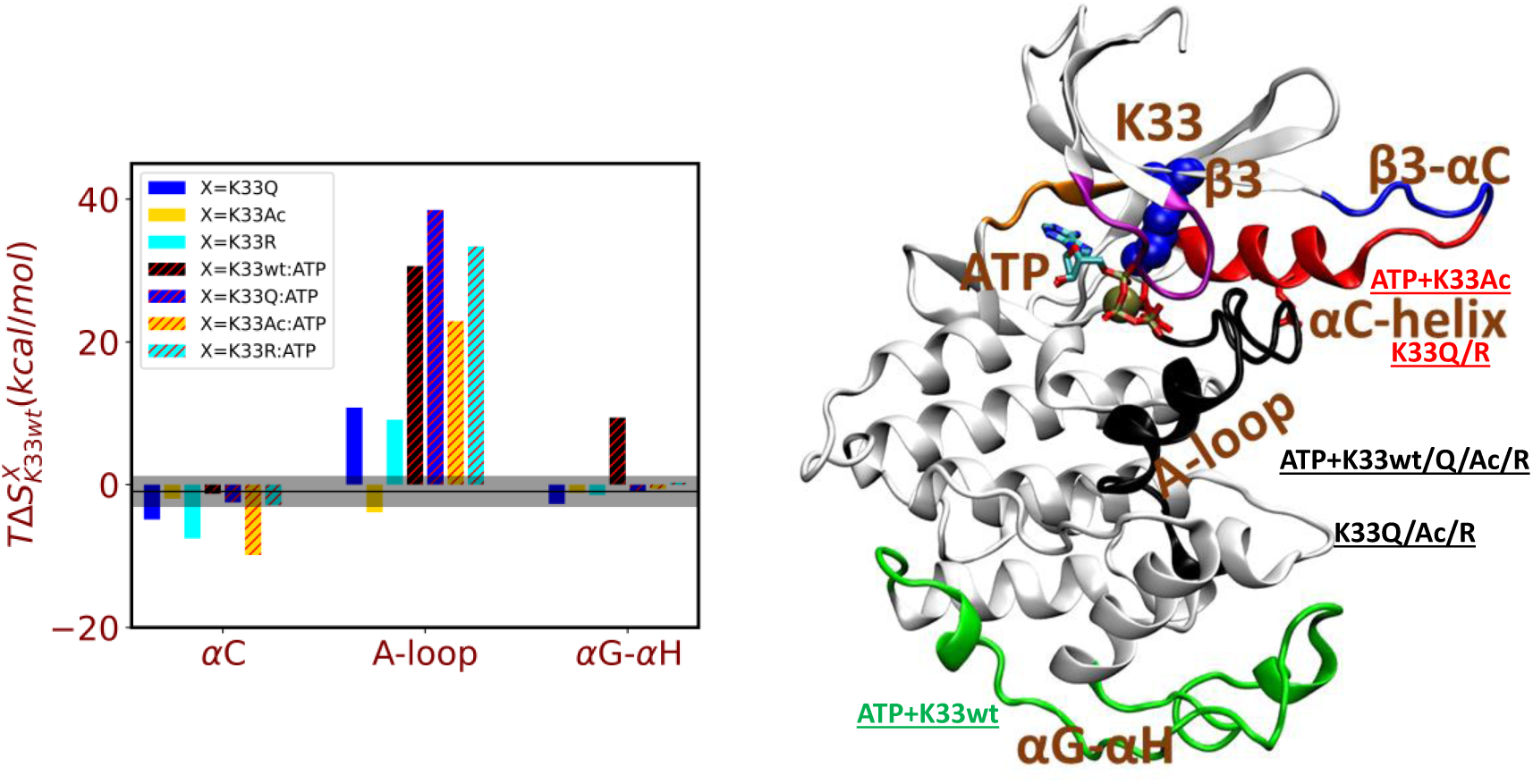
Summary of changes in the entropic component of free energy for K33wt upon ATP binding or mutations/acetylation at K33 (Left) Contributions to the changes in the entropic component 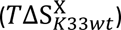 of free energy of the three PPIs in K33wt with active site perturbations (data from Fig. 2c and 3b). The horizontal black shaded region shows the baseline entropy changes across the protein. A value above this baseline is considered as a significant change in the entropic component of free energy. The centre of the baseline (black line) represents the average 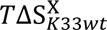 across all segments excluding the αC, A-loop, and αG-αH region. The width of the baseline (grey shaded region) is twice the standard deviation of the 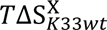 across all segments, excluding the αC, A-loop, and αG-αH region. The width is ∼ > 3.5 × thermal energy at *T*=303 *K* . **(Right)** Structure of CDK1 showing the relative positions of the three PPIs (red: 𝛼C-helix, black: A-loop, and green: 𝛼𝐺-𝛼H-loop) to the bound ATP (stick representation) and K33 (blue vdW spheres). We summarize the active site perturbations that have a significant effect on each PPI in the order of their significance (red, black, green text).

We have also rigorously examined the correlation between entropic changes and changes in thermal fluctuations in protein segments (**Fig. 4** and ESI **Fig. S10-12**) based on a large set of MD trajectories. Since the translational and rotational entropies do not change across the systems (**Table S6**), effectively the analysis examines the correlation between vibrational entropy and fluctuations of segments. We find that perturbing the K33wt active site creates entropy changes and variance changes in the same protein segments, the locally placed L7, 3, and G-loop and the remotely placed 𝛼𝐶-helix, A-loop, and 𝛼𝐺-𝛼H-loop. Averaging over the full set of trajectories, an unfavourable increase (decrease) in entropy of a protein segment following a perturbation is associated with a decrease in its flexibility as intuitively expected. However, we find that, in a significant subset of trajectory pairs, associated with perturbed/unperturbed states, the variance changes of protein segments do not correlate. These observations are puzzling as both the vibrational entropy and the variance of protein segments arise from the same variance-covariance matrix of atomic fluctuations (**subsection 2.3**). To better understand this issue, we decomposed the entropy and variance contributions from different regions of the protein further across all principal component modes for a pair of K33wt and K33wt:ATP trajectories (data shown in **Fig. 4c**) which exhibits a discrepancy between entropy and fluctuations (**Fig. S13-15**). Clearly, the entropy and the variance changes correlate when the first 2 slowest modes are included, however, as higher frequency modes are included, the correlation breaks down. This reflects a difference in the dependence of the entropy and variance changes on the mode frequencies. Whereas, the variance is dominated by contributions from the slowest modes, entropy contributions decay slowly with increasing mode frequency. We find that variance across segments changes stop evolving after including the top 5 lowest frequency modes, but entropy changes continue to evolve upon including higher frequency modes and seems to saturate only after including the top ∼300 modes (**Fig. S13-15**). Therefore, correlating protein dynamics with entropy based on a small statistical sample of MD trajectories or individual trajectory pairs might be misleading.

## 5. Conclusions

We have carried out a comprehensive computational thermodynamic and statistical analysis of active site perturbations (ATP binding, acetylation/mutations at K33) in CDK1. Our analysis reveals that ATP binding affinities are reduced for all mutants relative to K33wt. In all systems other than K33R, both enthalpy and entropy contribute significantly to determine the overall ATP-binding affinity. For K33R, ATP binding is dominated by enthalpic changes. ATP binding leads to both local and non-local entropy changes. Interestingly, the long range entropy contributions were found to favour ATP binding in all four systems. Projecting the mode entropies into contributions from protein segments revealed that in addition to local active site protein segments (L7, 3, and G-loop), the 𝛼𝐶-helix, 𝛼𝐺-𝛼𝐻 loop, and the A-loop, which interface with Cyclin, Cks and substrate proteins, regulate ATP binding and the stability of mutations/acetylation at K33. We find that while dynamical allosteric changes induced by active site perturbations are localized on the same protein segments as entropic changes, on average across a large set of MD trajectories. However, these quantities do not correlate in a significant fraction of the trajectories. In conclusion, our study reveals the mechanisms of ATP binding to free CDK1 and reveals allosteric communications (dynamic and entropic) between the enzyme active site and multiple PPIs which interface with Cyclin-B, substrate, and Cks proteins. Further studies examining the thermodynamics of Cyclin/Cks/substrate protein binding to CDK1, may reveal how these two distinct modes of allosteric couplings might be harnessed to selectively modulated PPIs by perturbations at the active site.

## Acknowledgment

K.V. and R.V. acknowledge funding support from the Department of Atomic Energy (DAE), Government of India, under project no. 12-R&D-TFR-5.10-0100 (TIFR 19P0153). U.K.-S. acknowledges support from the following funding sources: TIFR/DAE (19P0116 & ARUMDA 19P0911) and Department of Biotechnology (BT/PR29878/PFN/20/1431/2018).

## Notes

### Competing Interest Statement

The authors have declared no competing interest.

