## Supporting Data (Figures and Tables) for "ATP-Binding Free Energy Simulations Reveal an Allosteric Link Between the Enzyme Active Site and Multiple Functional Protein-Protein Interaction Interfaces in Cyclin Dependent Kinase-1"

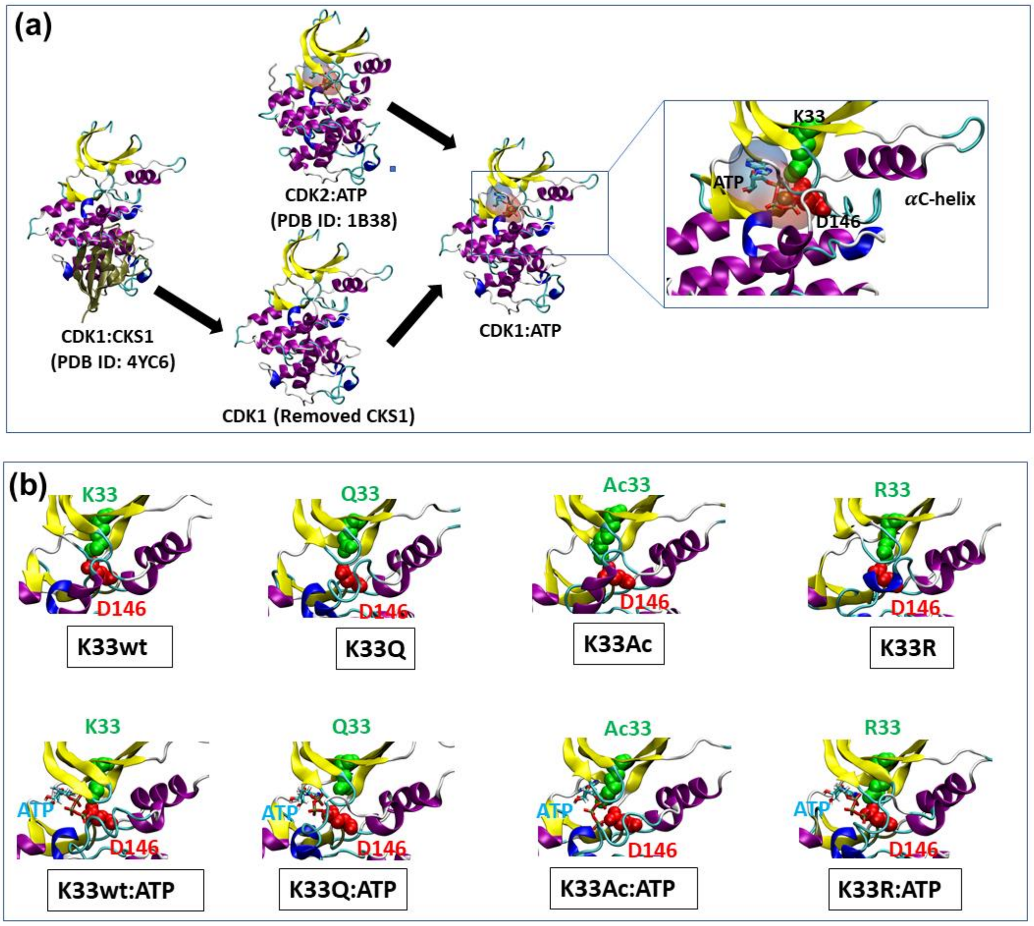


***Fig. S1.*** *Modeling of eight different monomeric states of CDK1.* ***(a)*** *Modelling of ATP bound CDK1. The CDK2 and CDK1 structures are superimposed to dock the ATP in the active site of CDK1 from CDK2.* ***(b)*** *The equilibrated active site geometry of each of the eight systems highlighting K/Q/Ac/R33 and D146 residues in VDW representation*

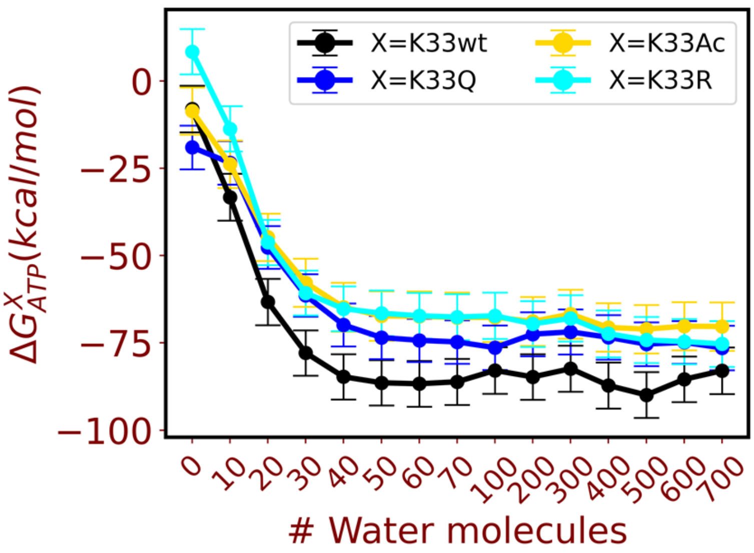


***Fig. S2.*** *Changes in free energy ($\Delta G_{ATP}^{X}$) associated with ATP binding as a function of n (from 0 to 700) number of most proximal water molecules to D146$C_\alpha$ and whose oxygen atoms are within 3.5 angstroms of any of protein's heavy atoms. We show mean values for these quantities and the standard error of mean extracted from 60×50 ns trajectories for each system.*

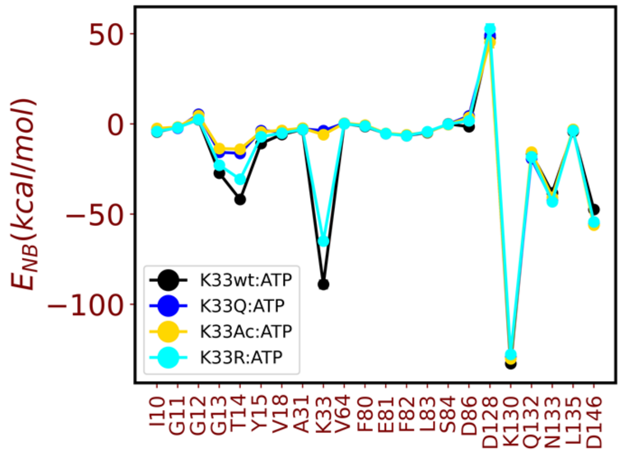


***Fig. S3.*** *Non-bonded interaction energies (Electrostatic + van der Waals) between ATP-Mg and its neighboring residues (residues with any heavy atom ≤4 Å away from ATP-Mg in crystal structure) in K33wt:ATP, K33Q:ATP, K33Ac:ATP, and K33R:ATP. We show mean values for these quantities and the standard error of mean extracted from 60×50 ns trajectories for each system.*

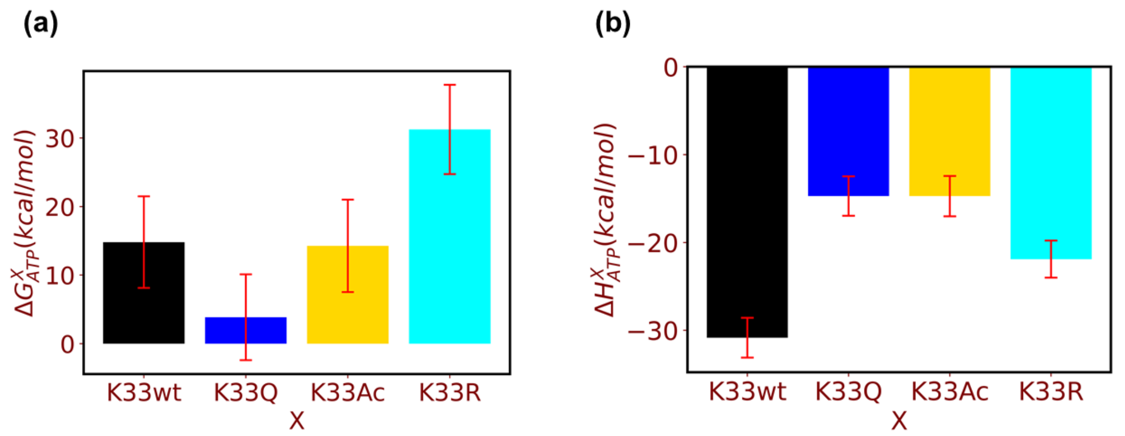


***Fig. S4. (a)*** *Mean free energy change* ($\Delta G^{X}_{ATP}$) *associated with ATP binding in the four systems K33wt, K33AC, K33Q and K33R excluding the contribution of active site water molecules. Here* $X$ *indicates* $K33wt/Q/Ac/R$. ***(b)*** *Corresponding mean enthalpy change* ($\Delta H^{X}_{ATP}$) *associated with ATP binding in the four systems K33wt, K33AC, K33Q and K33R excluding the contribution of active site water molecules. Here* $X$ *indicates* $K33wt/Q/Ac/R$. *We show mean values for these quantities and the standard error of mean extracted from 60×50 ns trajectories for each system.*

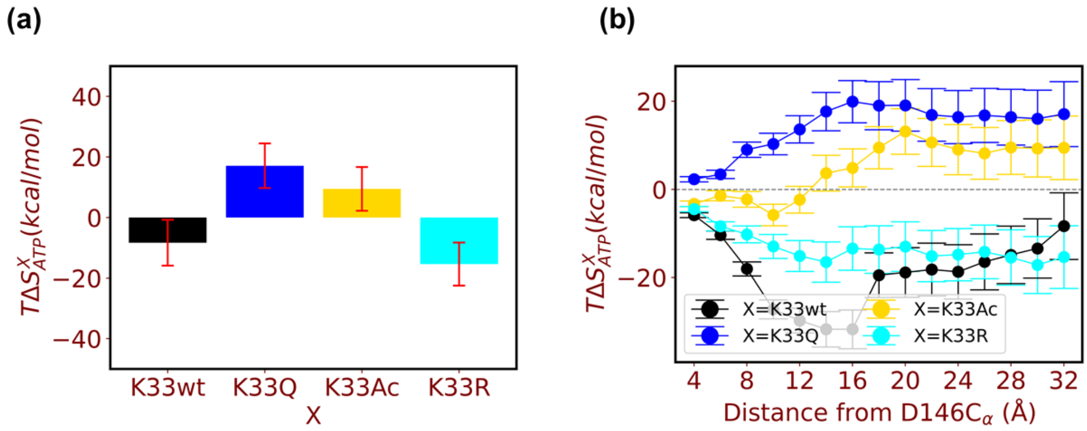


***Fig. S5***. ***(a)*** *Changes in vibrational entropy ($\Delta S^{X}_{ATP}$) of CDK1 associated with ATP binding in K33wt, K33AC, K33Q and K33R.* ***(b)*** *Changes in vibrational entropy ($\Delta S^{X}_{ATP}$) of CDK1 associated with ATP binding in K33wt, K33AC, K33Q and K33 . as a function of distance from the D146-Cα atoms . Here $X$ indicates $K33wt/Q/Ac/R$. Here, for a given distance d we compute entropy change associated with non-hydrogen atoms of residues lying within a sphere of radius d centered at the D146-Cα atom. For d=32 Å, all residues of the proteins are included. We show mean values for these quantities extracted from 60×50 ns trajectories for each system.*

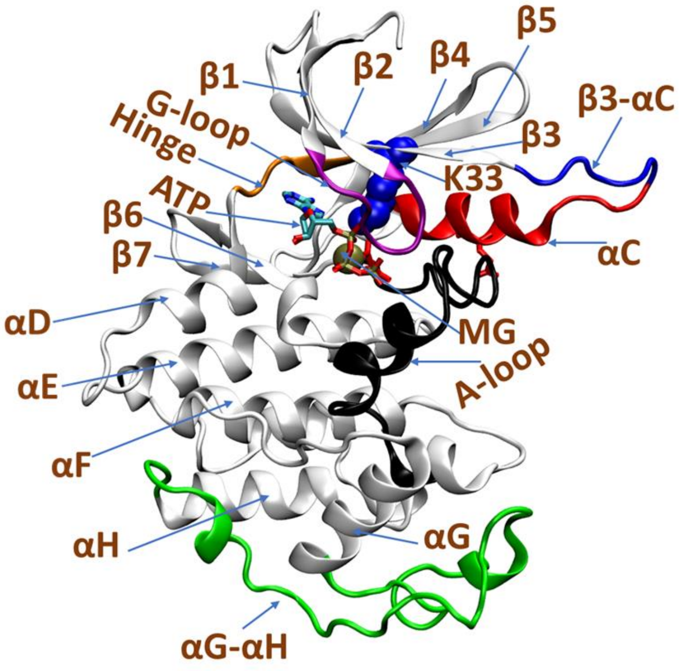


***Fig. S6.*** *Cartoon representation of CDK1 showing different segments for which entropy change in CDK1 are decomposed. The segments L1, L2… to L11 are not shown here. They correspond to the region between the segments shown in the above figure. Table S7 provides a list of residues contained in each segment.*

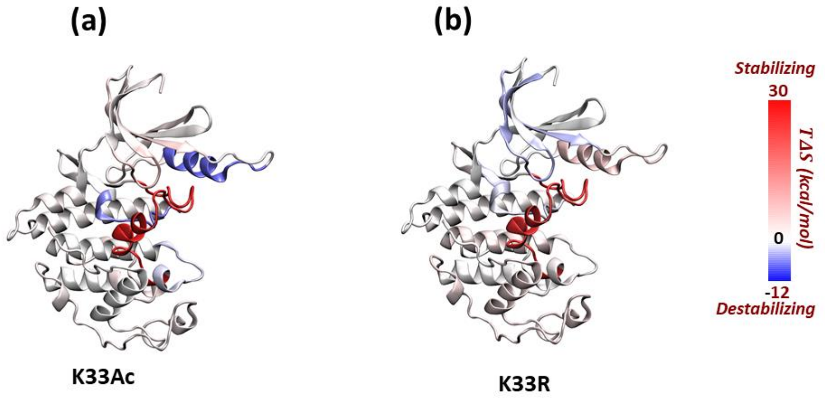


***Fig. S7.*** *Heat maps showing the entropy changes upon ATP binding in the protein structure for **(a)** K33Ac and **(b)** K33R.*

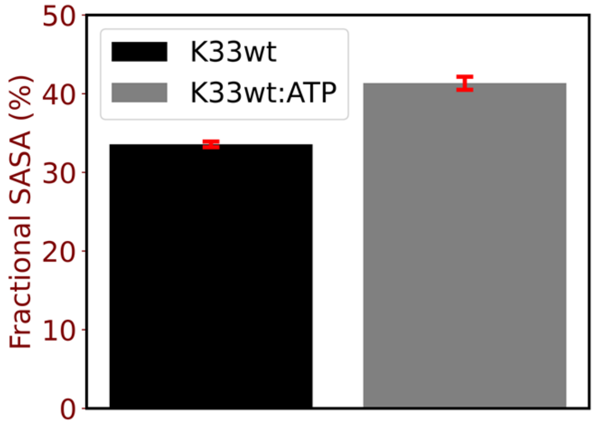


***Fig. S8.*** *Fractional solvent accessible surface area (SASA) of T161 in K33wt and K33wt:ATP. Fractional SASA is defined as the ratio between SASA of ATP in presence of surrounding residues to SASA in absence of surrounding residues multiplied by 100.*

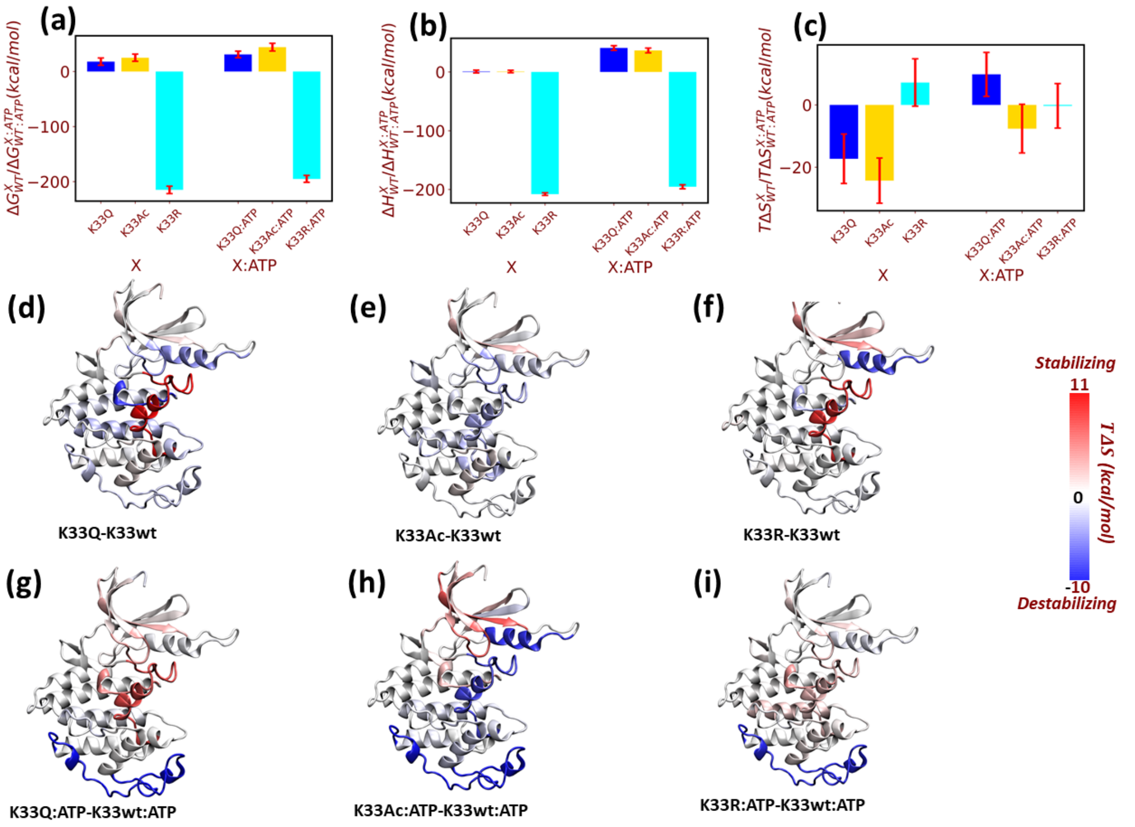


***Fig. S9.*** *Relative values of **(a)** free energy ($\Delta G^{X}_{K33wt/K33wt:ATP}$), **(b)** enthalpy ($\Delta H^{X}_{K33wt/K33wt:ATP}$), and **(c)** entropy ($\Delta S^{X}_{K33wt/K33wt:ATP}$) of K33Q/Ac/R and K33Q/Ac/R:ATP with respect to K33wt and K33WT:ATP, respectively. Here, averages and standard errors of the mean were extracted from 60×50 ns trajectories for each system (Here X=K33Ac, K33Q, and K33R for ATP-free CDK1 and X=K33Ac:ATP, K33Q:ATP, and K33R:ATP for ATP-bound CDK1). **(d-i)** Heat maps showing the entropy changes in the protein structure corresponding to (c).*

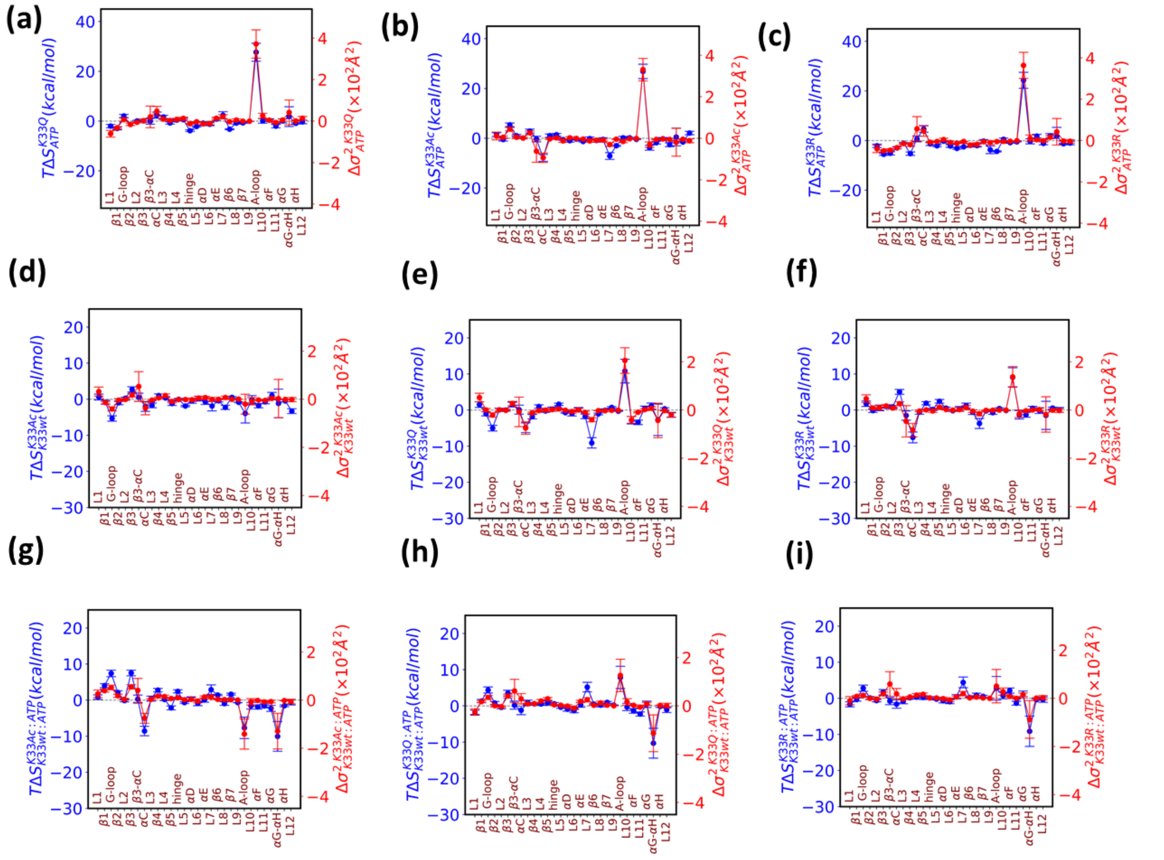

***Fig. S10**. Correlation between the variance change (red) and entropy change (red) upon **(a-c)** ATP binding in K33AC, K33Q, and K33R. **(d-f)** mutation in ATP-free **(g-i)** and ATP-bound CDK1 from the 30 non-overlapping segments of CDK1. Here, averages and standard errors of the mean were extracted from 60×50 ns trajectories for each system*

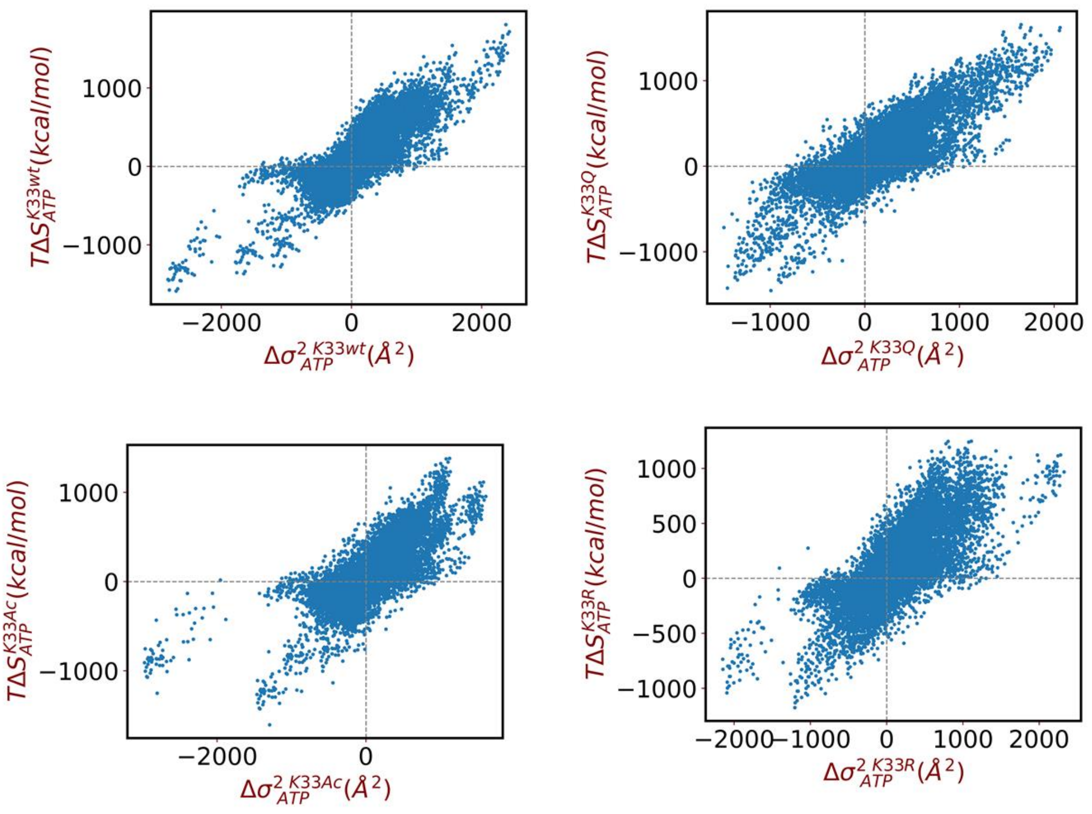


***Fig. S11.*** *Scatter plot depicting the relationship between variance change (x-axis) and entropy change (in energy unit, y-axis) upon ATP binding in K33wt, K33AC, K33Q, and K33R from 30 non-overlapping segments of CDK1. Each point on the plot represents the variance and entropy change from one of the non-overlapping segments 30 protein segments (refer to Table S7 for details) from a pair of trajectories, each obtained for ATP-free and ATP-bound states.*

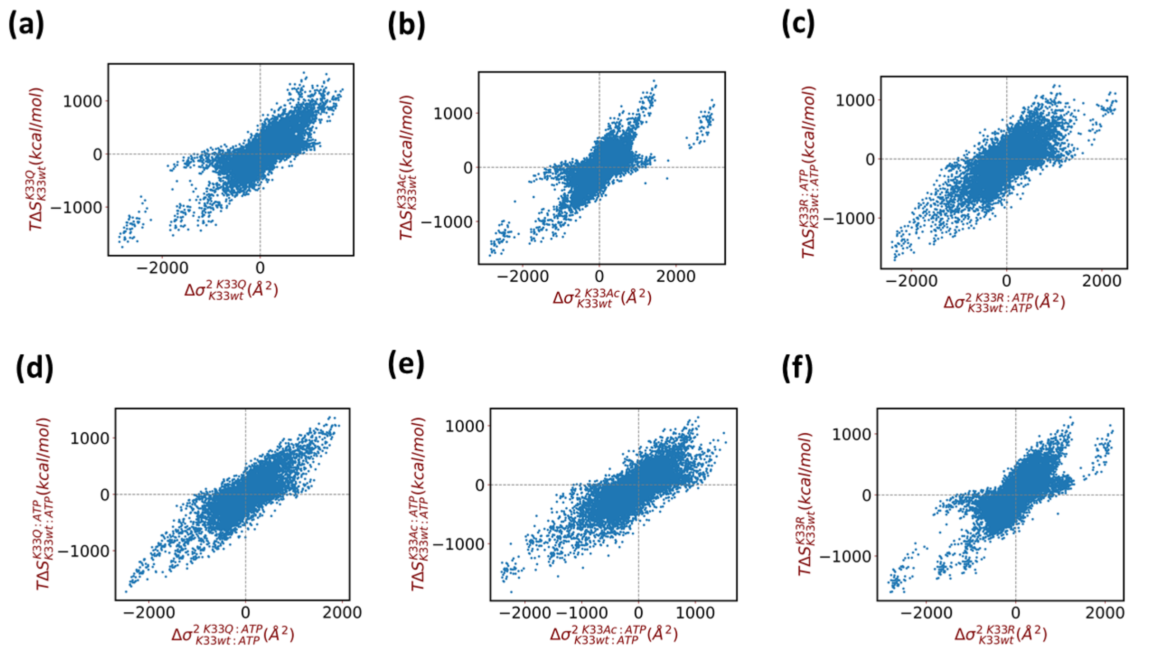


***Fig. S12.*** *Scatter plot depicting the relationship between variance change (x-axis) and entropy change (in energy unit, y-axis) upon mutations in ATP-free* ***(a-c)*** *and ATP-bound CDK1* ***(d-f)****, from 30 non-overlapping segments of CDK1. Each point on the plot represents the variance and entropy change from one of the 30 non-overlapping protein segments (refer to Table S7 for details) from a pair of trajectories, each obtained for ATP-free and ATP-bound states.*

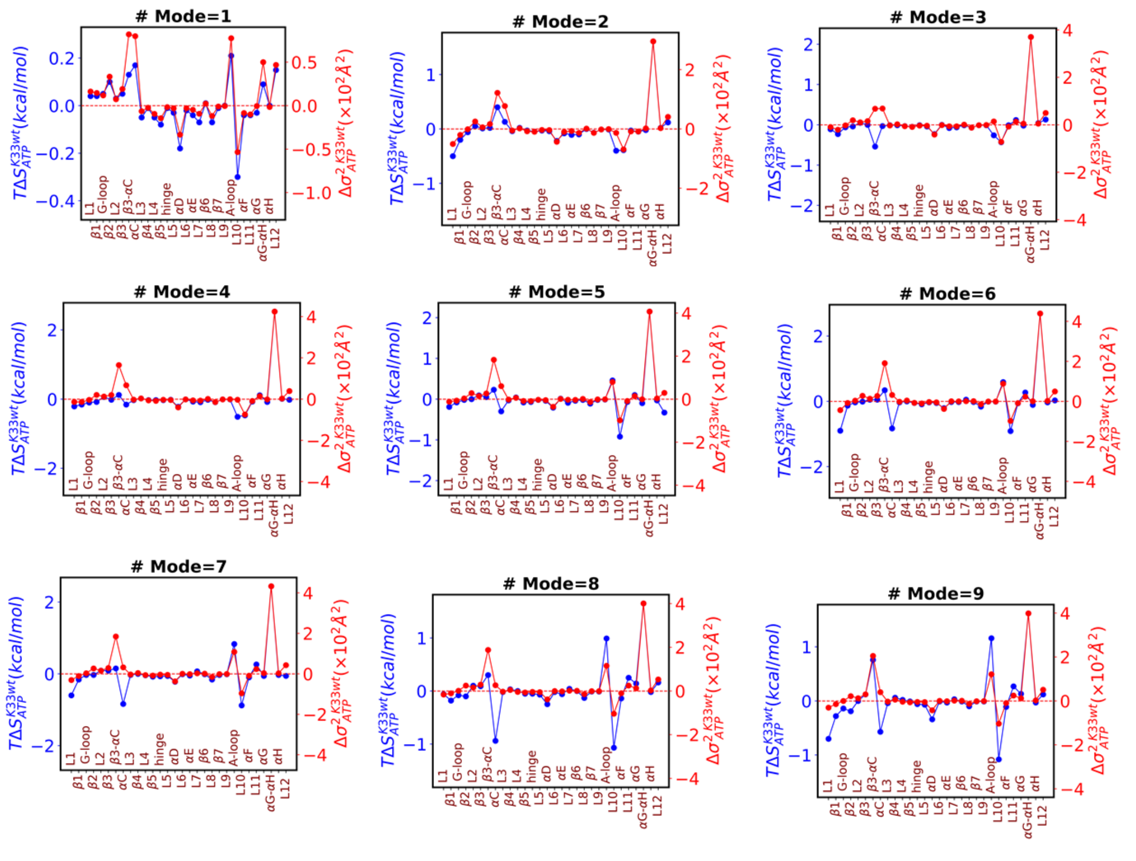


***Fig. S13.*** *Correlation between the variance and entropy change. Entropy contributions (blue) for different numbers of principal component modes to the free energies change upon ATP binding in K33wt from the 30 non-overlapping segments of CDK1 along with the corresponding variance changes (red), for a specific pair of trajectories corresponding to Fig. 4c. # Mode=X (X=1,2,..,9) indicates that first X slowest modes are included for entropy calculation.*

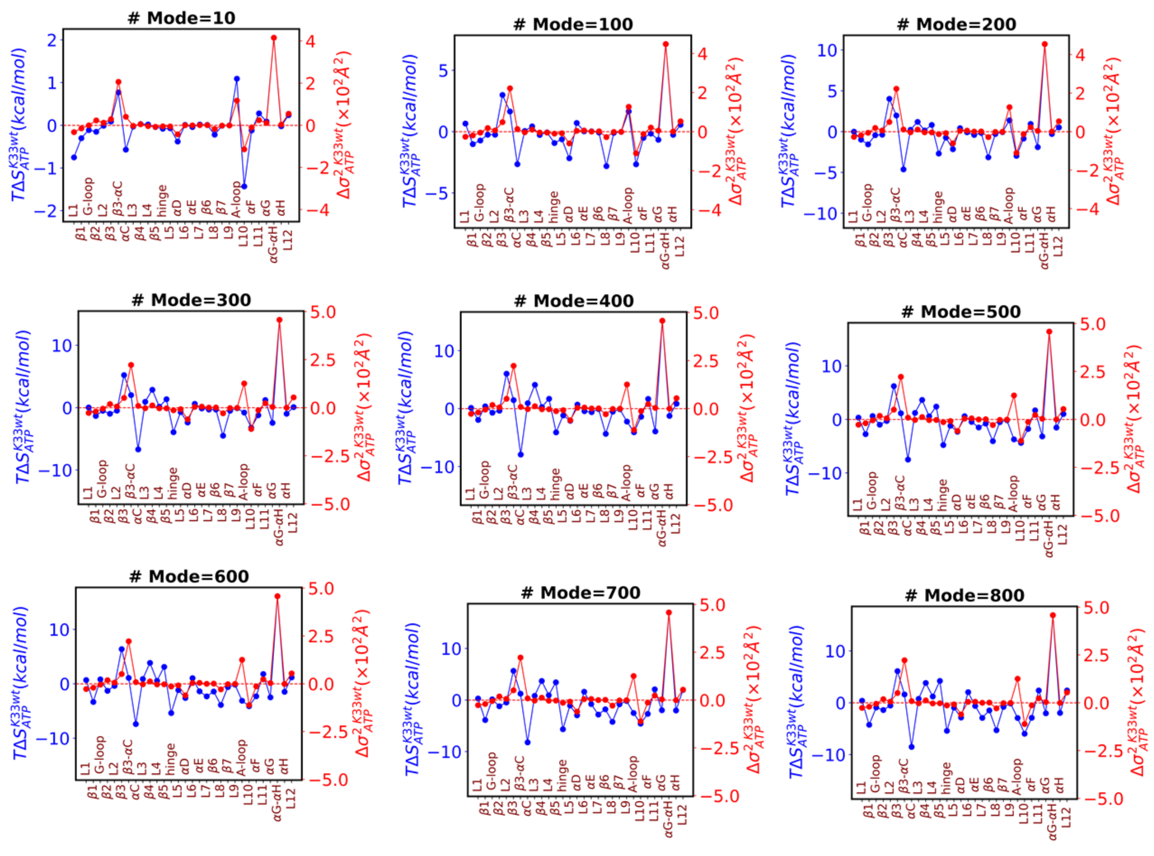


***Fig. S14.*** *Correlation between the variance and entropy change. Entropy contributions (blue) for different numbers of principal component modes to the free energies change upon ATP binding in K33wt from the 30 non-overlapping segments of CDK1 along with the corresponding variance changes (red), for a specific pair of trajectories corresponding to Fig. 4c. # Mode=X (X=10,100,200,300,400,500,600,700,800) indicates that first X slowest modes are included for entropy calculation.*

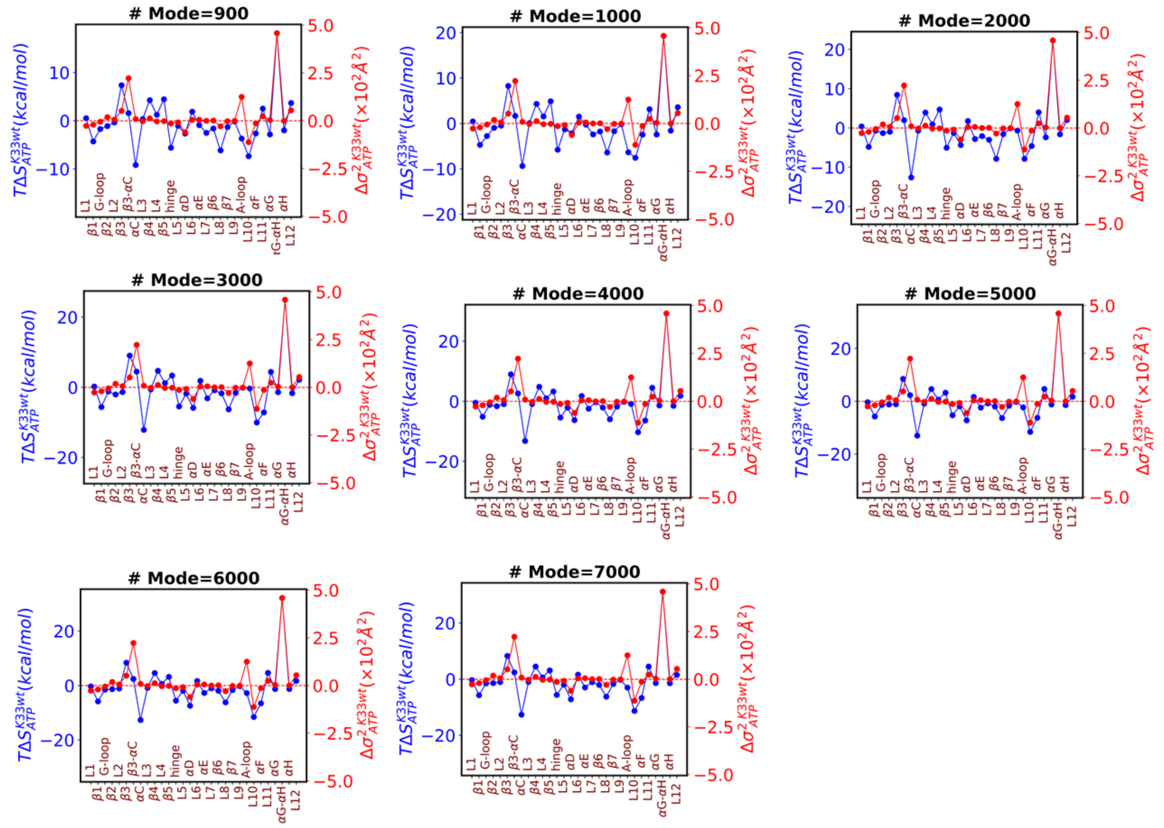

***Fig. S15.*** *Correlation between the variance and entropy change. Entropy contributions (blue) for different numbers of principal component modes to the free energies change upon ATP binding in K33wt from the 30 non-overlapping segments of CDK1 along with the corresponding variance changes (red), for a specific pair of trajectories corresponding to Fig. 4c. # Mode=X (X=900,1000,2000,3000,4000,5000,6000,7000) indicates that first X slowest modes are included for entropy calculation.*

| Models | Box Dimension (X, Y and Z) Å | Total Number of Atoms | Number of protein atoms | Number of Solvent Atoms | Number of Ions ($Na^+$\$Cl^-$) | Number of independent 50 ns trajectories |
|---|---|---|---|---|---|---|
| K33wt | (93x93x93) | 71997 | 4709 | 67287 | 1($Cl^-$) | 60 |
| K33Q | (93x93x93) | 71991 | 4704 | 67287 | 0 | 60 |
| K33Ac | (93x93x93) | 72000 | 4713 | 67287 | 0 | 60 |
| K33R | (93x93x93) | 71999 | 4711 | 67287 | 1($Cl^-$) | 60 |
| K33wt:ATP | (93x93x93) | 71912 | 4709 | 67158 | 1($Na^+$) | 60 |
| K33Q:ATP | (93x93x93) | 71911 | 4704 | 67161 | 2($Na^+$) | 60 |
| K33Ac:ATP | (93x93x93) | 71908 | 4713 | 67149 | 2($Na^+$) | 60 |
| K33R:ATP | (93x93x93) | 71914 | 4711 | 67158 | 1($Na^+$) | 60 |

***Table S1.*** *Values of simulation parameters (box dimension, total number of atoms, number of protein atoms, solvent atoms, ions, and number of MD trajectories) for all models.*

| Sampling Rate (ps) | Enthalpy | Error in enthalpy | Entropy | Error in entropy |
|---|---|---|---|---|
| 200 | -4489.6 | 10.6 | -5964.4 | 491.6 |
| 20 | -4490.3 | 10.5 | -4080.9 | 219.2 |
| 2 | -4490.8 | 10.5 | -4153.2 | 43.8 |

***Table S2.*** *Variation of average enthalpy and entropy with different sampling rates of MD frames in K33wt. Here enthalpy, entropy and error in their values, represents mean and error of the respective quantities extracted from 60×50 ns trajectories.*

| Energy terms | $\Delta G_{ATP}^{K33Q}/\Delta G_{ATP}^{K33wt}$ | $\Delta G_{ATP}^{K33Ac}/\Delta G_{ATP}^{K33wt}$ | $\Delta G_{ATP}^{K33R}/\Delta G_{ATP}^{K33wt}$ |
|---|---|---|---|
| bond | -0.352 | 0.187 | -0.357 |
| angle | -1.351 | 1.593 | -1.836 |
| dihedral | -5.204 | -7.333 | -3.001 |
| improper | 0.097 | -0.555 | -0.003 |
| electrostatic | 141.25 | 84.387 | 105.767 |
| vdW | 26.574 | 21.307 | 4.733 |
| solvation | -120.978 | -63.762 | -92.865 |

***Table S3.*** *Decomposition of enthalpy change of ATP binding in K33Q/Ac/R ($\Delta G_{ATP}^{K33Q}$, $\Delta G_{ATP}^{K33Ac}$, and $\Delta G_{ATP}^{K33R}$) relative to K33wt ($\Delta G_{ATP}^{K33wt}$) into different terms: bond, angle, dihedral, improper, electrostatic, vdW and solvation energy. Here, averages were extracted from 60×50 ns trajectories for each system. All energy values are in kcal/mol.*

.

| Energy terms | $\Delta G_{ATP}^{K33Q}/\Delta G_{ATP}^{K33wt}$ | $\Delta G_{ATP}^{K33Ac}/\Delta G_{ATP}^{K33wt}$ | $\Delta G_{ATP}^{K33R}/\Delta G_{ATP}^{K33wt}$ |
|---|---|---|---|
| bond | -0.142 | 0.62 | 0.011 |
| angle | -0.896 | 1.893 | -1.923 |
| dihedral | -5.388 | -7.222 | -3.162 |
| improper | 0.308 | -0.341 | 0.156 |
| electrostatic | 142.89 | 68.87 | 108.671 |
| vdW | 24.952 | 24.048 | 3.996 |
| solvation | -145.608 | -71.753 | -98.806 |

***Table S4.*** *Decomposition of enthalpy change of ATP binding in K33Q/Ac/R ($\Delta G_{ATP}^{K33Q}$, $\Delta G_{ATP}^{K33Ac}$, and $\Delta G_{ATP}^{K33R}$) relative to K33wt ($\Delta G_{ATP}^{K33wt}$) into different terms: bond, angle, dihedral, improper, electrostatic, vdW and solvation energy. During enthalpy calculation explicit water molecules at the active site are not taken into account. Here, averages were extracted from 60×50 ns trajectories for each system. All energy values are in kcal/mol.*

**(a)**

| | With active site water molecules | | | Without active site water molecules | | |
|---|---|---|---|---|---|---|
| **Energy terms** | $\Delta H_{K33wt}^{K33Q}$ | $\Delta H_{K33wt}^{K33Ac}$ | $\Delta H_{K33wt}^{K33R}$ | $\Delta H_{K33wt}^{K33Q}$ | $\Delta H_{K33wt}^{K33Ac}$ | $\Delta H_{K33wt}^{K33R}$ |
| bond | 0.249 | 1.297 | 4.252 | -0.194 | 0.654 | 3.673 |
| angle | -0.661 | 1.2 | 4.021 | -0.91 | 1.133 | 4.455 |
| dihedral | 5.817 | 1.136 | 2.646 | 5.573 | 0.484 | 2.282 |
| improper | 0.407 | 0.689 | 0.079 | 0.24 | 0.619 | 0.07 |
| electrostatic | -42.567 | -56.56 | -286.702 | -48.527 | -58.47 | -285.281 |
| vdW | -13.276 | -11.089 | -1.302 | -9.3 | -8.827 | -0.377 |
| solvation | 50.858 | 64.173 | 69.198 | 58.172 | 65.187 | 72.481 |

**(b)**

| | With active site water molecules | | | Without active site water molecules | | |
|---|---|---|---|---|---|---|
| **Energy terms** | $\Delta H_{K33wt:ATP}^{K33Q:ATP}$ | $\Delta H_{K33wt:ATP}^{K33Ac:ATP}$ | $\Delta H_{K33wt:ATP}^{K33R:ATP}$ | $\Delta H_{K33wt:ATP}^{K33Q:ATP}$ | $\Delta H_{K33wt:ATP}^{K33Ac:ATP}$ | $\Delta H_{K33wt:ATP}^{K33R:ATP}$ |
| bond | -0.103 | 1.484 | 3.895 | -0.336 | 1.273 | 3.684 |
| angle | -2.012 | 2.793 | 2.185 | -1.806 | 3.026 | 2.532 |
| dihedral | 0.612 | -6.196 | -0.355 | 0.185 | -6.738 | -0.88 |
| improper | 0.503 | 0.134 | 0.076 | 0.548 | 0.279 | 0.226 |
| electrostatic | 98.683 | 27.827 | -180.935 | 94.364 | 10.4 | -176.61 |
| vdW | 13.298 | 10.219 | 3.43 | 15.652 | 15.22 | 3.619 |
| solvation | -70.12 | 0.411 | -23.667 | -87.436 | -6.567 | -26.326 |

***Table S5.*** *Decomposition of enthalpy change **(a)** between K33Q/Ac/R and K33wt ($\Delta H_{K33wt}^{K33Q}$, $\Delta H_{K33wt}^{K33Ac}$, and $\Delta H_{K33wt}^{K33R}$) and **(b)** between K33Q/Ac/R:ATP and K33wt:ATP ($\Delta H_{K33wt:ATP}^{K33Q:ATP}$, $\Delta H_{K33wt:ATP}^{K33Ac:ATP}$, and $\Delta H_{K33wt:ATP}^{K33R:ATP}$) into different terms: bond, angle, dihedral, improper, electrostatic, vdW and solvation, both in presence (left panel) and absence (right panel) of explicit active site water molecules. Here, averages were extracted from 60×50 ns trajectories for each system. All energy values are in kcal/mol.*

| X | $\Delta G^{X}_{ATP}$ | $\Delta H^{X}_{ATP}$ | $T\Delta S^{X}_{ATP}$ (vibration) | $T\Delta S^{X}_{ATP}$ (rotation) | $T\Delta S^{X}_{ATP}$ (translation) |
|---|---|---|---|---|---|
| K33wt | -63.6 ± 6.5 | -109.2 ± 2.8 | -22.8 ± 7.6 | -11.3 | -11.5 |
| K33Q | -50.6 ± 6.2 | -69.2 ± 3.6 | 4.3 ± 7.4 | -11.3 | -11.5 |
| K33Ac | -44.4 ± 7.1 | -73.4 ± 3.5 | -6.1 ± 7.5 | -11.3 | -11.5 |
| K33R | -43.7 ± 6.5 | -96.8 ± 3.0 | -30.3 ± 7.2 | -11.3 | -11.5 |

***Table S6.*** *Decomposition of free energy change upon ATP binding in K33wt, K33Q, K33Ac, and K33R ($\Delta G^{K33wt}_{ATP}$, $\Delta G^{K33Q}_{ATP}$ $\Delta G^{K33Ac}_{ATP}$, and $\Delta G^{K33R}_{ATP}$). Here we have shown changes in free energy (2nd column), enthalpy (3rd column), vibrational entropy (4th column), rotational entropy (5th column), translation entropy (6th column). The temperature T corresponds to 303 K, same as temperature used in MD simulation. All energy values are in kcal/mol.*

| Segment Name | Residue Range | Segment Name | Residue Range |
|---|---|---|---|
| L1 | 1-3 | L6 | 97-101 |
| $\beta 1$ | 4-10 | $\alpha$E | 102-123 |
| G-loop | 11-17 | L7 | 124-132 |
| $\beta 2$ | 18-24 | $\beta 6$ | 133-136 |
| L2 | 25-26 | L8 | 137-141 |
| $\beta 3$ | 27-37 | $\beta 7$ | 142-144 |
| $\beta 3$-$\alpha$C | 38-43 | L9 | 145-145 |
| $\alpha$C | 44-57 | A-loop | 146-173 |
| L3 | 58-65 | L10 | 174-183 |
| $\beta 4$ | 66-71 | $\alpha$F | 184-199 |
| L4 | 72-73 | L11 | 200-205 |
| $\beta 5$ | 74-79 | $\alpha$G | 206-220 |
| hinge | 80-84 | $\alpha$G-$\alpha$H | 221-257 |
| L5 | 85-87 | $\alpha$H | 258-268 |
| $\alpha$D | 88-96 | L12 | 269-289 |

***Table S7.*** *Segment name and corresponding residue range for which changes in entropy in CDK1 are decomposed. Residue range for each segments besides L1-L20, are taken from the reference 1.*

| Segment Name | # Pair of trajectories | | | | | | | | | % of Pair of trajectories showing non-correlation |
|---|---|---|---|---|---|---|---|---|---|---|
| | $\Delta\sigma^2(+)\Delta S(+)$ | $\Delta\sigma^2(+)\Delta S(0)$ | $\Delta\sigma^2(+)\Delta S(-)$ | $\Delta\sigma^2(0)\Delta S(+)$ | $\Delta\sigma^2(0)\Delta S(0)$ | $\Delta\sigma^2(0)\Delta S(-)$ | $\Delta\sigma^2(-)\Delta S(+)$ | $\Delta\sigma^2(-)\Delta S(0)$ | $\Delta\sigma^2(-)\Delta S(-)$ | |
| $G$-loop | 233 | 106 | 52 | 63 | 154 | 280 | 13 | 107 | 2592 | 17 |
| $\beta 3$ | 491 | 261 | 345 | 233 | 188 | 379 | 221 | 271 | 1211 | 48 |
| $\alpha C$ | 1000 | 138 | 160 | 98 | 41 | 68 | 305 | 265 | 1525 | 29 |
| $L7$ | 310 | 97 | 110 | 44 | 75 | 388 | 8 | 32 | 2536 | 19 |
| $A$-loop | 3061 | 34 | 50 | 45 | 7 | 8 | 146 | 38 | 211 | 9 |
| $\alpha G$-$\alpha H$ | 1767 | 75 | 198 | 81 | 11 | 46 | 324 | 87 | 1011 | 22 |

***Table S8.*** *Number trajectory pairs (ATP-bound vs ATP-free CDK1) for correlations observed between $\Delta\sigma^2$ and $\Delta S$ in specific segments upon ATP binding in K33wt. For each segment, we have 3600 pair of trajectories (from 60x50ns trajectories each for K33wt and K33wt:ATP). We count the number of pairs for each type of correlation observed: positive $\Delta\sigma^2$ and positive $\Delta S$ ($\Delta\sigma^2(+)\Delta S(+)$), positive $\Delta\sigma^2$ and zero change in $\Delta S$ ($\Delta\sigma^2(+)\Delta S(0)$), positive $\Delta\sigma^2$ and negative $\Delta S$ ($\Delta\sigma^2(+)\Delta S(-)$), zero change in $\Delta\sigma^2$ and positive $\Delta S$ ($\Delta\sigma^2(0)\Delta S(+)$), zero change in $\Delta\sigma^2$ and zero change in $\Delta S$ ($\Delta\sigma^2(0)\Delta S(0)$), zero change in $\Delta\sigma^2$ and negative $\Delta S$ ( $\Delta\sigma^2(0)\Delta S(-)$), negative $\Delta\sigma^2$ and positive $\Delta S$ ($\Delta\sigma^2(-)\Delta S(+)$), negative $\Delta\sigma^2$ and zero change in $\Delta S$ ($\Delta\sigma^2(-)\Delta S(0)$), negative $\Delta\sigma^2$ and negative $\Delta S$ ($\Delta\sigma^2(-)\Delta S(-)$). The criteria for positive, zero and negative change in $\Delta S$ and $\Delta\sigma^2$ are established by assessing the standard deviations $\delta\Delta S_{other}$ and $\delta\Delta\sigma^2{}_{other}$ in entropy change and variance change across all segments excluding the G-loop, $\beta 3$, $\alpha C$, L7, A-loop, and $\alpha$G-$\alpha$H. The criteria for positive, zero and negative change in $\Delta S$ are as follows: $+\Delta S$ if $\Delta S > \delta\Delta S_{other}$, $-\Delta S$ if $\Delta S < -\delta\Delta S_{other}$ $0$ $\Delta S$ if $-\delta\Delta S_{other} <= \Delta S <= \delta\Delta S_{other}$. Similarly the criteria for positive, zero and negative change in $\Delta\sigma^2$ are: $+\Delta\sigma^2$ if $\Delta\sigma^2 > \delta\Delta\sigma^2{}_{other}$, $-\Delta\sigma^2$ if $\Delta\sigma^2 < -\delta\Delta\sigma^2{}_{other}$, and $0$ $\Delta\sigma^2$ if $-\delta\Delta\sigma^2{}_{other} \leq \Delta\sigma^2 <= \delta\Delta\sigma^2{}_{other}$.*
